# Speed tuning in head coordinates as an alternative explanation of depth selectivity from motion parallax in area MT

**DOI:** 10.1101/2021.04.17.440243

**Authors:** Zhe-Xin Xu, Gregory C. DeAngelis

## Abstract

There are two distinct sources of retinal image motion: motion of objects in the world and movement of the observer. In cases where an object moves in a scene and the eyes also move, a coordinate transformation that involves smooth eye movements and retinal motion will be needed in order to estimate object motion in world coordinates. More recently, interactions between retinal and eye velocity signals have also been suggested to generate depth selectivity from motion parallax (MP) in the macaque middle temporal (MT) area. We explored whether the nature of the interaction between eye and retinal velocities in MT neurons favors one of these two possibilities or a mixture of both. We analyzed responses of MT neurons to retinal and eye velocities in a viewing context in which the observer translates laterally while maintaining visual fixation on a world-fixed target. In this scenario, the depth of an object can be inferred from the ratio between retinal velocity and eye velocity, according to the motion-pursuit law. Previous studies have shown that MT responses to retinal motion are gain-modulated by the direction of eye movement, suggesting a potential mechanism for depth tuning from MP. However, our analysis of the joint tuning profile for retinal and eye velocities reveals that some MT neurons show a partial coordinate transformation toward head coordinates. We formalized a series of computational models to predict neural spike trains as well as selectivity for depth, and we used factorial model comparisons to quantify the relative importance of each model component. Our findings for many MT neurons reveal that the data are equally well explained by gain modulation or a partial coordinate transformation toward head coordinates, although some responses can only be well fit by the coordinate transform model. Our results highlight the potential role of MT neurons in representing multiple higher-level sensory variables, including depth from MP and object motion in the world.

## Introduction

Living in a dynamic, three-dimensional world, many animals rely heavily on visual information to infer the motion and depth of objects. The retinal image motion of an object generally reflects both self-motion (including translation and rotation of the head, as well as rotation of the eyes) and movements of objects in the world. Depending on the viewing context, retinal motion could serve as a powerful cue for different properties of an object, such as its depth or its velocity relative to the world.

When an object is stationary in the world and a moving observer fixates a world-fixed target, retinal velocity depends on the distance from the object to the observer’s point of fixation (Howard & Rogers, 2002). In this case, the retinal image motion (or motion parallax, MP) provides crucial information about the depth of the object. Theoretical and psychophysical studies have shown that depth perception from MP depends on the ratio of retinal velocity to eye velocity, as specified by the motion-pursuit law (Nawrot & Stroyan, 2009). Moreover, previous studies show that humans utilize these signals to judge depth (Bradshaw & Rogers, 1996; Nawrot, 2003; Ono et al., 1988; Ono et al., 1986; Rogers & Graham, 1979, 1982; Rogers, 1993). Neurophysiological studies have demonstrated that neurons in the middle temporal (MT) area of macaque monkeys are selective for the sign (near/far) of depth defined by MP (Nadler et al., 2008), and that this selectivity is realized by combining retinal image motion with smooth eye movement command signals (Nadler et al., 2009). A more recent study reported that MT responses are gain modulated by the direction of eye movement to generate depth-sign selectivity (Kim et al., 2017). Moreover, previous work has also shown that MT neurons are selective for depth from MP when eye rotation is simulated by large-field visual motion, such that “dynamic perspective” cues are provided (Kim et al., 2015). These findings together suggest that signals related to smooth eye movements in area MT may serve the computational goal of extracting depth from MP. So far, however, there are no direct measurements of how MT neurons are jointly tuned for eye velocity and retinal velocity, which would clarify the exact nature of the interaction between these variables in the neural responses.

When an object is moving in the world and a stationary observer tracks the moving target with their eyes, the retinal velocity of the object is the difference between eye velocity and the object’s velocity relative to the scene. To estimate the object’s velocity in head or world coordinates, the visual system must subtract the image motion produced by eye movement. This might require an interaction between retinal and eye velocity signals that could be different from the interaction involved in computing depth from MP, but this issue has not been examined. Previous studies have examined how neurons in areas MT and MST represent object motion during smooth pursuit eye movements (Inaba et al., 2007, 2011; Chukoskie & Movshon, 2009). They measured neural speed tuning in both retinal and screen coordinates during pursuit eye movements, and investigated whether tuning in areas MT and MST was shifted toward screen coordinates when smooth eye movements were executed. While these studies generally found much greater effects in MST neurons, the response shifts of some MT neurons were not negligible. Thus, whether or not MT neurons convey useful information about object motion in the world remains an open question. Moreover, it remains unclear whether there is any relationship between selectivity for velocity in screen coordinates and selectivity for depth from MP.

It has long been established that neurons in area MT are sensitive to binocular disparity and local two-dimensional motion signals (Albright, 1984; Born & Bradley, 2005; DeAngelis & Newsome, 1999; Felleman & Kaas, 1984; Maunsell & Van Essen, 1983a; Maunsell & Van Essen, 1983b; Rodman & Albright, 1987; Zeki, 1974), and that their responses associate with the perception of depth and motion (Britten et al., 1996; Britten et al., 1992; DeAngelis et al., 1998; DeAngelis & Newsome, 2004; Uka & DeAngelis, 2004). However, relatively little is known about how MT neurons contribute to the computation of higher-level perceptual variables, such as depth from MP and object motion in the world. Existing evidence, as described above, suggests a potential role of MT neurons that goes beyond representing retinal motion, but the detailed mechanisms by which other variables, such as eye movements, modulate MT responses are not well understood.

In this study, we reanalyze existing data from experiments on depth from MP, and devise a novel way to closely examine the interactions between eye velocity and retinal image velocity in the responses of MT neurons. We formulate a model in which eye velocity modulates the gain of MT responses, substantially extending the work of Kim et al. (2017). We fit this model to MT spike trains and we show that the joint tuning for eye and retinal velocities is well described by this gain modulation model for many MT neurons. However, the joint tuning for other neurons reveals a clear diagonal structure that cannot be explained by a mechanism of gain modulation (Kim et al., 2017). We further develop a computational model to account for responses that are shifted, in varying degrees, toward head-centered speed tuning (i.e., tuning for object motion in the world). Our head-centered tuning model also successfully predicts the depth tuning curves of many MT neurons, including those with diagonal structure, and predicts an empirically-observed relationship between depth-sign selectivity and speed preference that was previously unexplained. By directly comparing these two classes of models, we assess the possibility that depth from MP could be jointly represented with head-centered object motion in area MT. Our findings raise the possibility that eye velocity signals in MT might be involved in partially shifting the representation of motion toward head coordinates, and call into question whether the main goal of eye velocity modulations in area MT is to compute depth from MP. Our simulations make specific testable predictions for experiments that could distinguish the two classes of models.

## Materials and Methods

### Neural recording data

Electrophysiological data examined in this study were taken from a series of studies on motion parallax in macaque area MT, which have been described in detail previously (Kim et al., 2015; Nadler et al., 2008; Nadler et al., 2009). In brief, extracellular single-unit responses were recorded using tungsten microelectrodes (FHC Inc). For each neuron, basic tuning properties including speed and direction tuning were measured, followed by a main experimental protocol that involved three stimulus conditions: 1) In the motion parallax (MP) condition (Figure 1A), a motion platform was used to laterally translate the animal sinusoidally along the preferred-null axis of each recorded neuron (see Nadler et al. 2008 for details). The stimulus was a 0.5 Hz sinusoid multiplied by a windowing function to avoid sharp onset acceleration, and had two starting phases (0 and 180 deg). A world-fixed target was presented throughout the movement, and animals were trained to fixate this target by making smooth pursuit eye movements in the opposite direction to head translation. A random-dot patch was placed over the neuron’s receptive field and viewed monocularly, with the motion of the patch computed to provide motion parallax consistent with one of nine simulated depths (0°, ±0.5°, ±1.0°, ±1.5°, ±2.0°). 2) In the retinal motion (RM) condition (Figure 1B), the animal remained stationary while fixating on a static target. The same image motion over the receptive field as in the MP condition was generated by moving the OpenGL camera in the virtual environment along the same trajectory as in the MP condition. Consequently, this condition contained identical image motion of the patch relative to the fixation target as presented in the MP condition, but no pursuit eye movements were made. The size of the random-dot patch was small enough that no significant information for eye movement was present in this condition (Kim et al. 2015). 3) In the dynamic perspective (DP) condition (Figure 1C), the image motion of the random-dot patch and the fixation target were the same as those in the RM condition. However, a 3D background of random dots was presented, whose image motion simulated optic flow that resulting from the body translation and compensatory smooth pursuit eye movements made in the MP condition (see Kim et al. 2015 for details). Thus, the background dots provided robust information about eye rotation relative to the scene in the absence of real eye movements.

**Figure 1.**
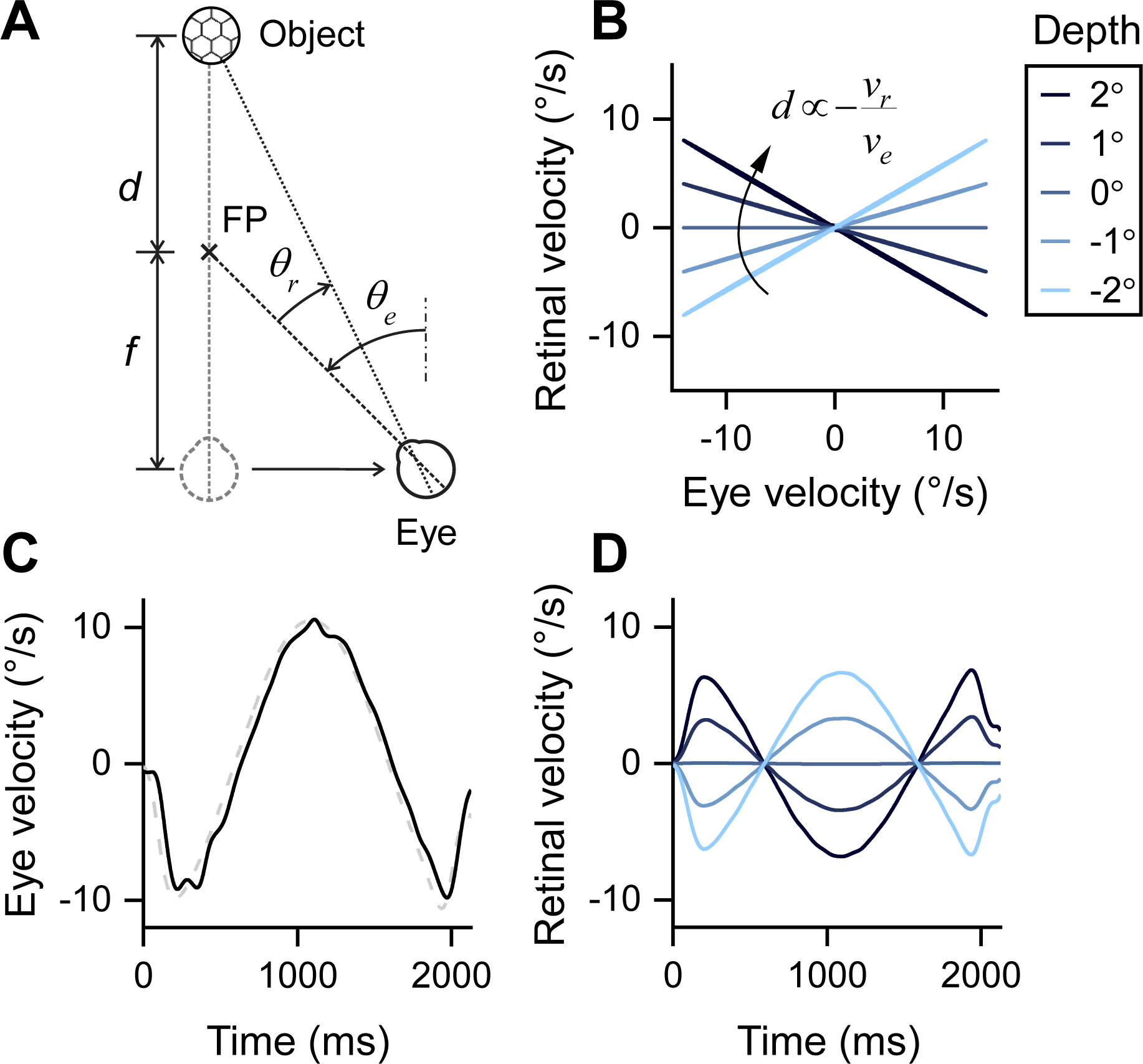
Retinal and eye velocities in motion parallax. **A**, Geometry of motion parallax. An observer is translating rightwards while tracking a stationary target *FP* at distance *f*. The relative depth, *d/f,* of an object can be obtained from its retinal image motion *dθ_r_ dt* = *v_r_* and the observer’s eye velocity, *dθ_e_ dt* = *v_e_*. **B**, Depths in retinal-eye velocity plane. In the 2D space of retinal and eye velocities, depth is described by the slope of each line that passes through the origin. Depths that are closer than the fixation point (“near”) have positive slopes (lighter blue lines), while far depths have negative slopes (darker blue lines). **C**, Temporal profile of averaged eye velocity in the experiment. **D**, Temporal trajectories of retinal velocities in different depths.

**Figure 2.**
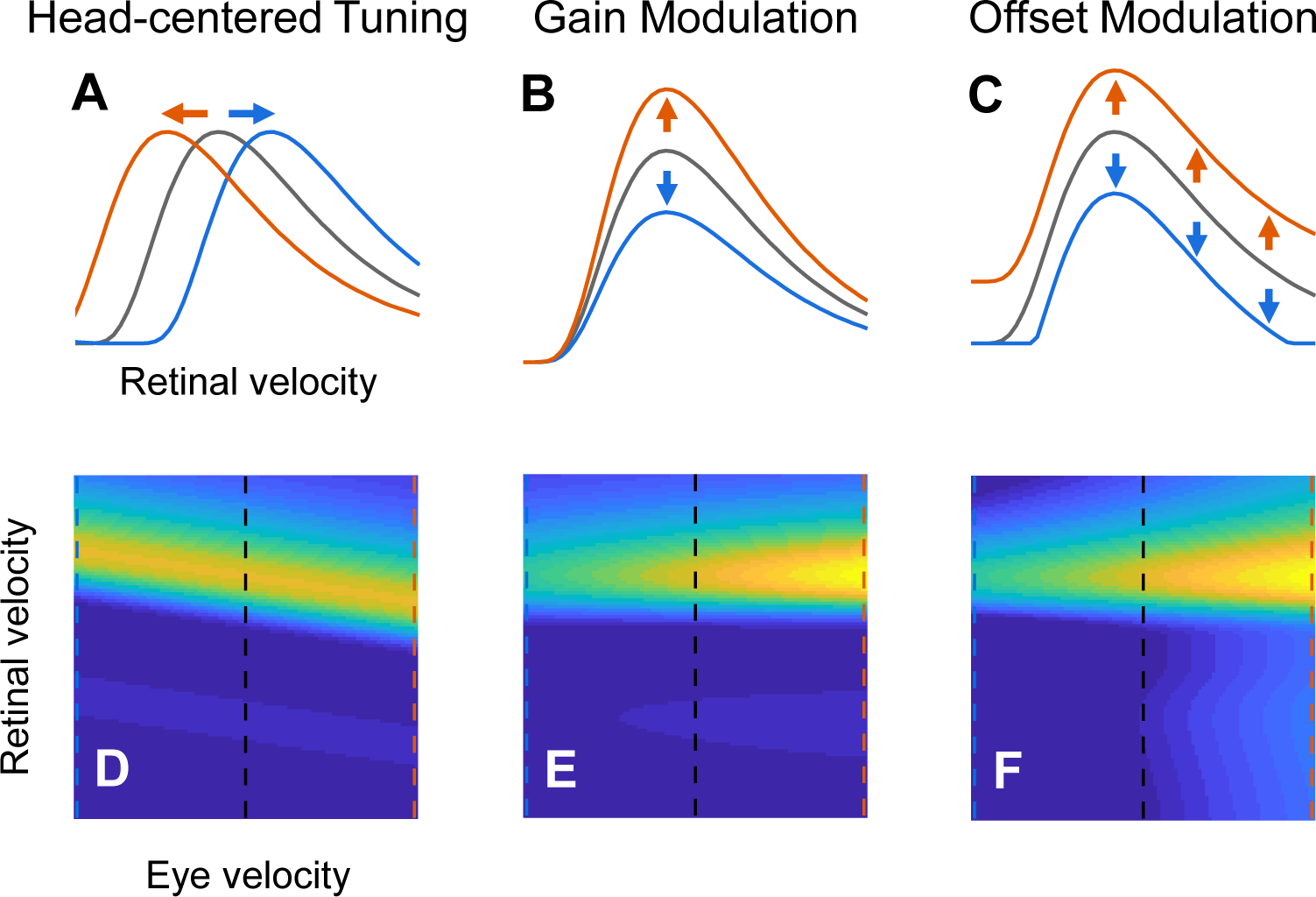
Possible interactions between retinal and eye velocities. **A**, Speed tuning that shifts toward head-centered coordinates. **B**, Multiplicative gain modulation on the retinal speed tuning curve. **C**, Additive offset modulation on the speed tuning curve. **D**-**F**, Joint tuning corresponding to each mechanism.

We excluded cells that did not show strong response modulations by retinal image motion in the following way. We first calculated the difference in firing rates between the two opposite movement phases (0° and 180°) for each depth in the RM condition. This subtraction canceled out baseline activity and stimulus onset transients that were not dependent on depth, allowing us to isolate the response modulation induced by image motion. A Fourier transform was then applied to obtain the amplitude spectrum. We took the median amplitude at frequencies in the range from 0.5 to 1 Hz (where the dominant energy in our stimulus resides), and computed the significance of that amplitude by shuffling spike trains 10,000 times across time bins. The significance level was Bonferroni corrected for multiple comparisons. Cells that showed significant modulation amplitudes at less than two depths with the same sign (both near or both far) were excluded from analyses. Approximately 8% of the cells were removed in this fashion, and most were neurons tuned to high speeds that were not well driven by our stimuli (which had a maximal retinal velocity around 7 °/s).

### Joint tuning for retinal and eye velocities

Vertical and horizontal eye position signals were recorded by an eye coil system at a sampling rate of 200 Hz. We linearly interpolated the raw data to 1 kHz, then rotated it to obtain signals aligned with the preferred-null axis of the neuron under study. The position data was filtered by a Gaussian window having *σ* = 33 ms. MATLAB function *filtfilt* was used to prevent phase shifts. The velocity of eye movements was then computed by taking the derivative of the position signals. We calculated the theoretical retinal image position of the center of the random-dot patch at each frame based on the simulated geometry and OpenGL projective parameters. We then calibrated this signal by adding the positional difference between the eye and target during pursuit to find the actual retinal image position of the center of the patch at each time point. The first derivative of a Gaussian filter (*σ* = 33 ms) was then applied to the image positions to obtain retinal stimulus velocities.

To estimate the joint tuning of MT neurons for retinal and eye velocities, we discretized the joint instantaneous velocities in each trial into 1×1°/s bins, and we counted the number of spikes associated with each pair of velocities (Figure 3). The mean firing rate in each 1×1°/s bin was calculated as the total number of spikes divided by the number of samples of this particular pair of retinal and eye velocities, and then multiplied by the 1 kHz sampling rate. Bins with less than 200 velocity samples were discarded from the analysis. In the MP condition, the actual eye velocity at each time point in the trial was used. In contrast, for the RM and DP conditions, the simulated eye velocity (i.e., the idealized eye velocity that would be consistent with the visual images presented) was used to plot the joint velocity tuning profiles.

**Figure 3.**
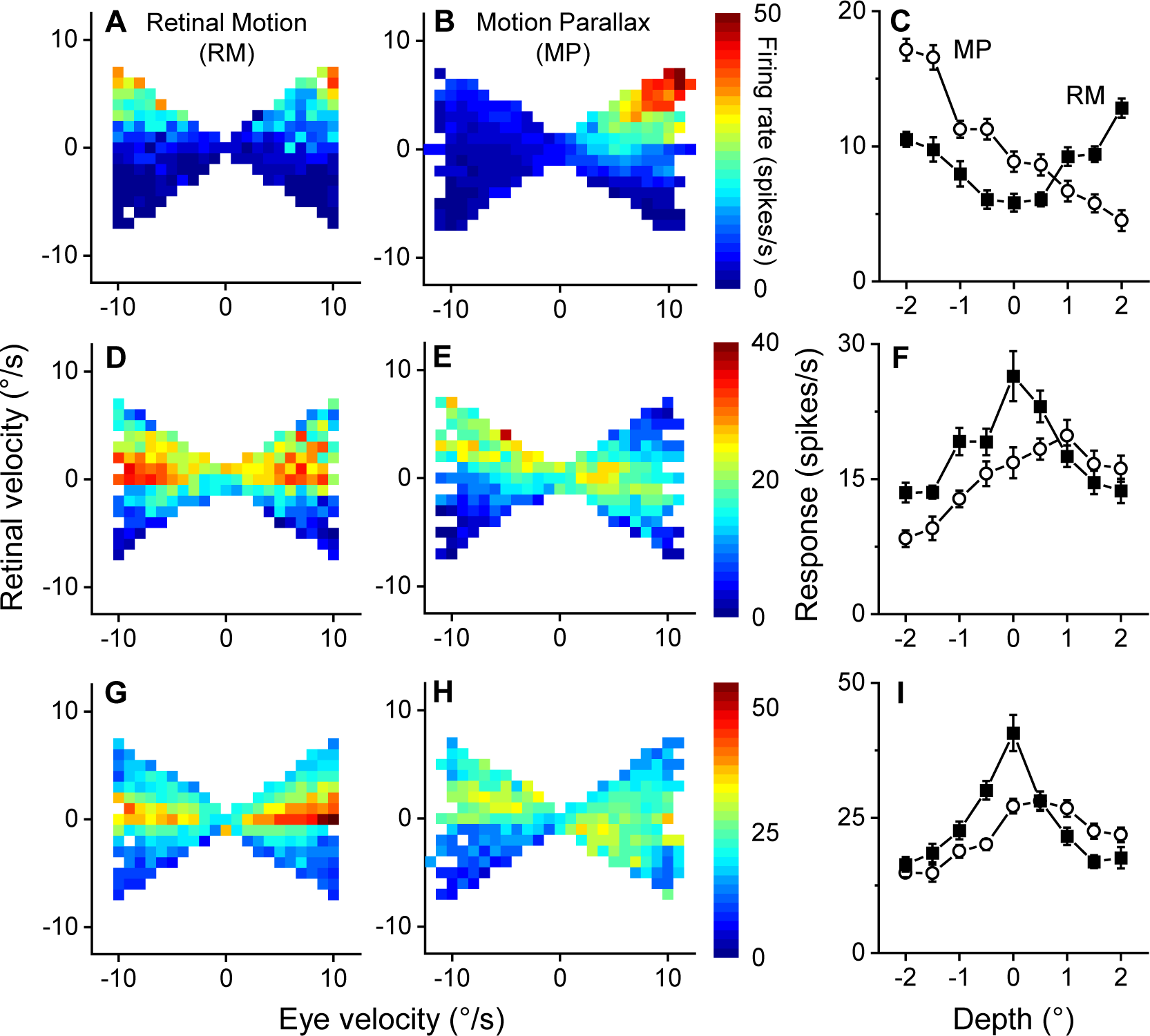
Examples of joint tuning and depth tuning curves. Each row corresponds to one example neuron. **A**, Joint tuning for retinal and eye velocities in the RM condition. **B**, Joint tuning in the MP condition. The response to retinal motion is suppressed by negative eye velocity (upper left part), and facilitated by positive eye velocity (upper right part). **C**, Depth tuning curve in both RM (solid squares) and MP (open circles) conditions. In the RM condition, depth tuning curve is symmetric around zero, showing no depth-sign preference. In the MP condition, the firing rate is higher for near depths and lower for far depths. **D** and **G**, Joint tuning in the RM condition for two example neurons with slow speed preference. **E** and **H**, Diagonal structure in the corresponding joint tuning in the MP condition. Responses around zero retinal velocity are not suppressed or enhanced by the eye velocity, but rather shift toward a negative diagonal direction. **F** and **I**, Depth tuning curves for far-preferring neurons. The depth tuning in the RM condition has a symmetric pattern with a peak at zero depth (solid squares). In the MP condition, the diagonal shift of joint tuning causes a far preference in depth tuning curves (open circles). Error bars in this figure indicate 1 SD of the mean.

### Simulation of head-centered speed tuning

To explore how depth-sign selectivity might arise for neurons that are tuned to velocity in head coordinates, we simulated neural responses with head-centered tuning and examined their joint tuning for retinal velocity and eye velocity (Fig. 5B), as well as their corresponding depth tuning curves. We simulated 500 neurons with velocity tuning that could range from being in retinal to head-centered coordinates, and with a diverse range of speed preferences. Velocity coordinates can be specified by a weighted sum between retinal and eye velocities, with an adjustable weight on eye velocity. For the *i*-th simulated neuron, spike rates ***r****_i_* sampled from a Poisson distribution:

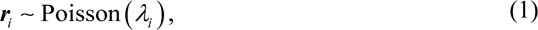

were randomly where *λ_i_* is the mean firing rate for a given pairing of retinal velocity ***v****_r_* and eye velocity ***v****_e_* :

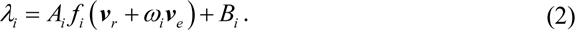

**Figure 4.**
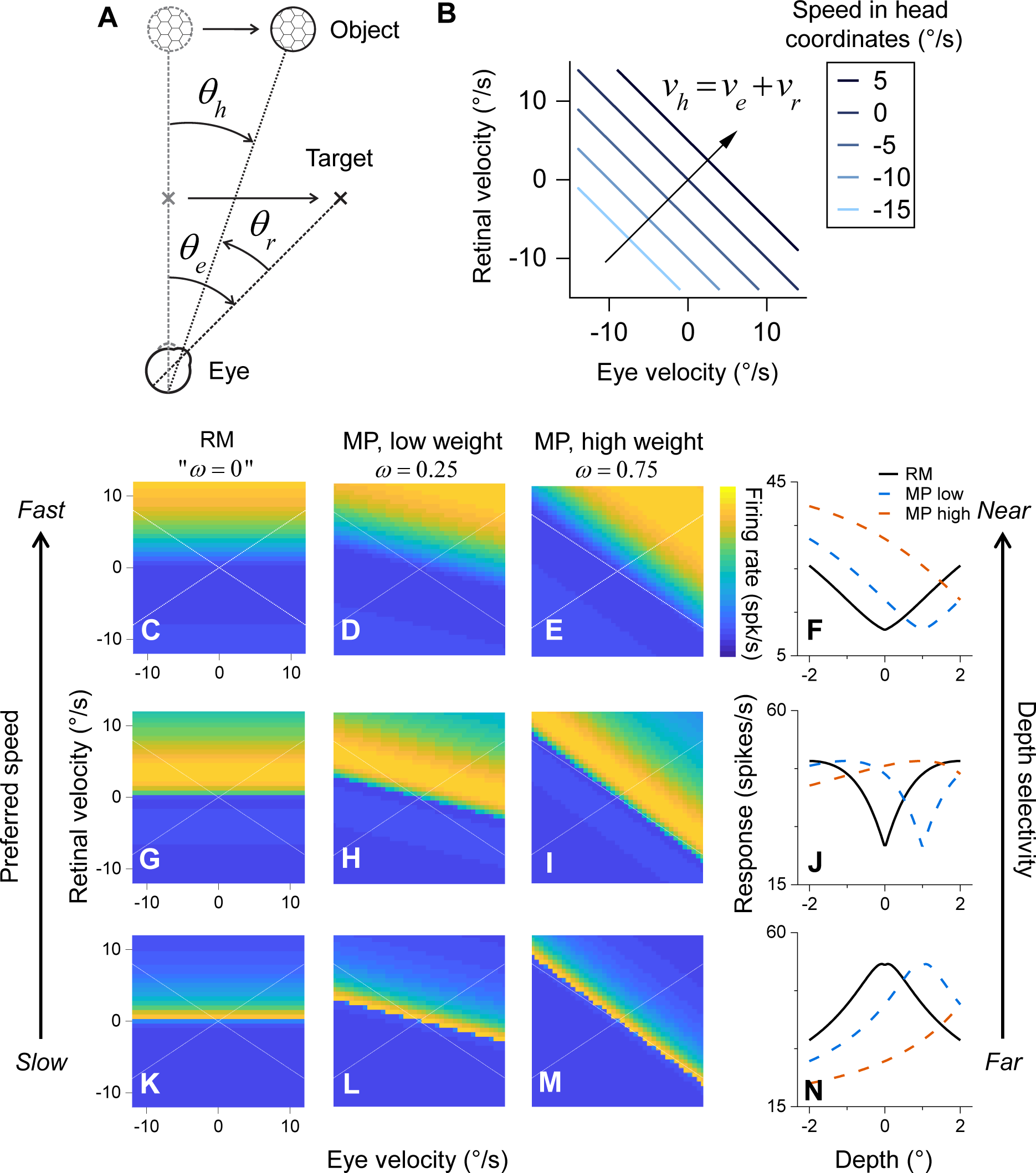
Velocity tuning in head coordinates. **A**, Geometry of head-centered object motion. While a stationary observer is tracking a moving target by smooth pursuit eye movements, an object also moves rightwards. The motion of the object in head-centered coordinates, *dθ_h_* / *dt* = *v_h_*, is the vector sum of its retinal motion *dθ_r_ dt* = *v_r_* and the observer’s eye velocity *dθ_e_ dt* = *v_e_*. **B**, Head-centered speed in retinal-eye velocity space. Each diagonal line indicates a particular speed in head coordinates. **C**, The joint tuning in RM condition for a simulated neuron with preferred speed at 15°/s. White dash lines show the range of samples in the experiment. **D** and **E**, Joint tuning in MP condition for low (*ω* = 0.25) and high (*ω* = 0.75) weights on eye velocity. **F**, Depth tuning curves in RM and MP conditions. Selectivity for depth occurs in the MP condition for both low and high weights (blue and orange dash lines), but not in the RM condition (solid black line). **G-J**, Similar with **A-E**, joint tuning and depth tuning curves for a model neuron with moderate preferred speed (3.3°/s). Depth selectivity again emerges in both weights of head coordinates in the MP condition. **K**-**N**, Similar with **A-E**, tuning patterns for a slow speed (0.16°/s) preferring neuron. A clear diagonal shift of the tuning response presents in both low (**L**) and high (**M**) weights, yielding a far-preferring depth tuning (**N**, dash lines).

**Figure 5.**
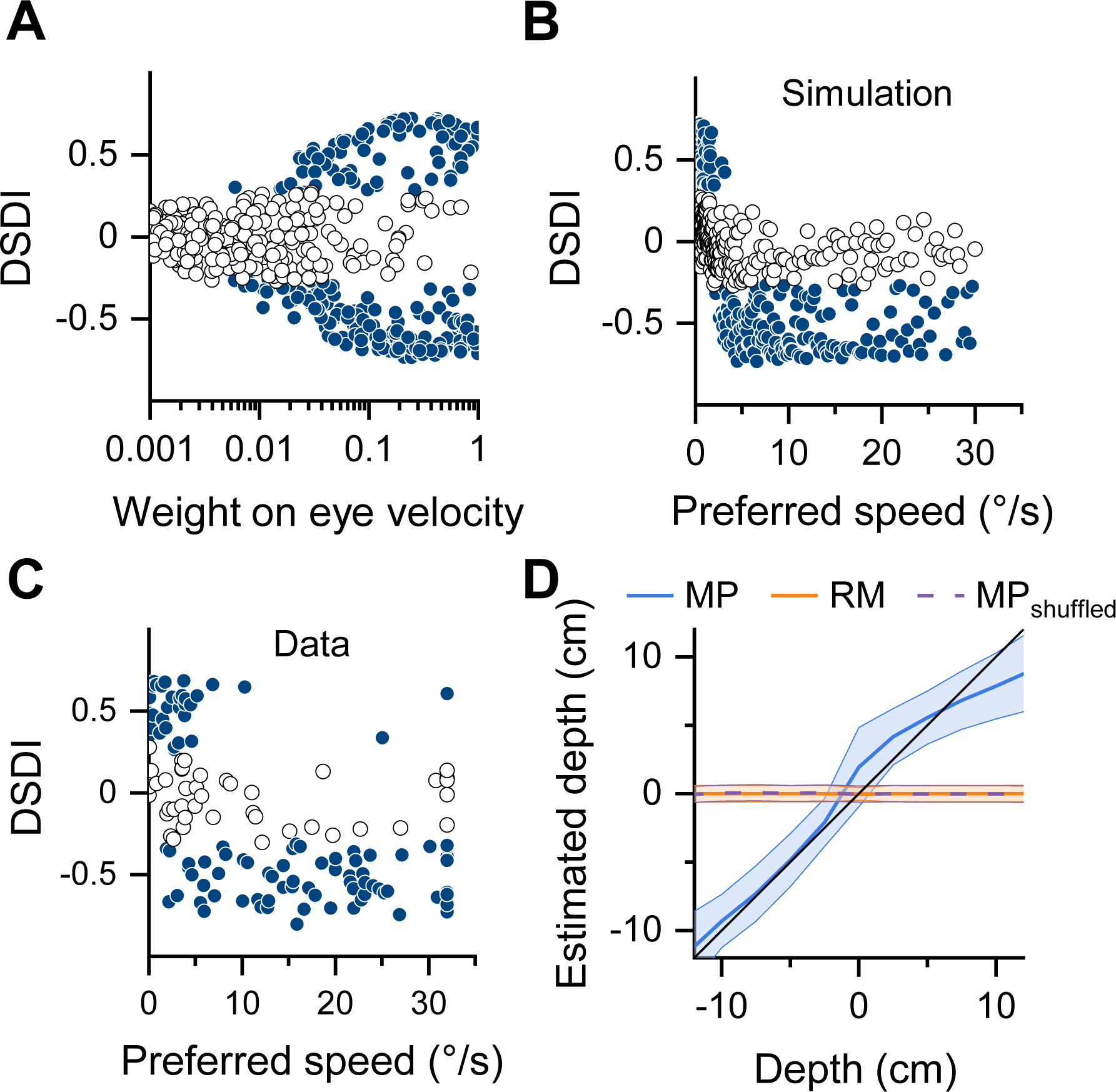
Population results of head-centered tuning simulation. **A**, Relationship between simulated depth-sign selectivity (DSDI) and the weight on eye velocity. Significant DSDI (*p* < .05, permutation test, solid dark blue circles) starts emerging when the weight on eye velocity is around 0.01, and saturates around 0.2. Open circles indicate non-significant DSDIs. **B**, Simulated DSDI is negatively correlated with preferred speed in head coordinates (Spearman’s *r* = -0.75, *p* < .001). **C**, Similar negative correlation between DSDIs and preferred speed in real data reported by Nadler et al. (2008) (Spearman’s *r* = -0.56, *p* < .001). **D**, Decoding performance of simulated population responses in RM and MP conditions. The decoder successfully estimated depth from MP responses (close to the identity line, blue curve), but failed in RM condition (flat orange curve). Shaded areas indicate 1 SD around the mean.

Here, *A_i_* is a scaling factor on response amplitude (uniformly sampled from 50 to 100 spikes/s), *B_i_* is the spontaneous firing rate (uniformly sampled from 0 to 20 spikes/s), and *ω_i_* is the weight placed on eye velocity (uniformly sampled in a logarithmic space of [0.001, 1]). *f_i_* (·) represents the velocity response function for each model neuron. It is well-established that the speed and direction tuning curves of MT neurons are multiplicatively separable (Rodman & Albright, 1987), and can be expressed as log-Gaussian (Nover et al., 2005) and von Mises functions (Smolyanskaya et al., 2013), respectively. Therefore, we modeled *f_i_* (·) as the product of these two functions:

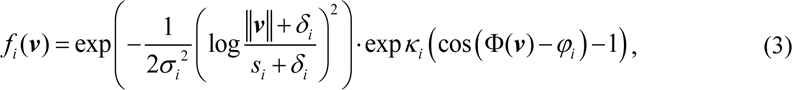

where ||***v***|| and Φ(***v***) indicate the magnitude (speed) and direction of a given 2D velocity vector 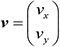 respectively. *s_i_* and *φ_i_* denote the preferred speed and direction, respectively, where *s_i_* is uniformly sampled in a logarithmic space from [0.1, 30] °/s. Because both retinal image motion and eye movements were aligned with each neuron’s preferred-null motion axis in each experiment, there is no need to simulate neurons with different direction preferences; we therefore simplified the model such that the preferred direction of each neuron was *φ_i_* = 0°. *σ_i_* and *κ_i_* determine the width of speed and direction tuning curves, respectively, with *σ _i_* ∈[0.5,1.5] and *κ_i_* ∈[1, 2]. *δ_i_* is a small constant that keeps the logarithm from being undefined as the speed ||***v***|| goes to zero, and was drawn from *δ_i_* ∈[0.001, 3]. All parameter ranges were chosen based on empirical data from area MT (Nover et al., 2005).

Given the fact that velocity in head coordinates is the sum of retinal and eye velocities, ***v****_h_* ≡ ***v****_r_* + ***v****_e_*, we can rewrite the input velocity of *f*(·) in Equation 2 for *ω_i_* ∈[0,1] :

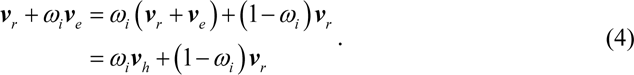

Hence, *ω_i_* is also the weight on head coordinates; the input velocity becomes completely retinocentric when *ωi* = 0, and purely head-centered when *ω_i_* = 1.

To obtain the joint velocity tuning and depth tuning for each simulated neuron, we presented pairs of retinal and eye velocities in a range of [-12, 12] °/s with 0.1°/s spacing. The direction of both velocities was along each neuron’s preferred-null axis (i.e., 0°). For each given depth, *d*, the depth tuning curve was computed as the averaged firing rate <***r****_i_*> across all stimulus combinations that are selected based on the motion-pursuit law (Nawrot & Stroyan, 2009):

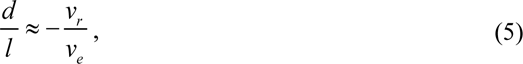

where *l* is the viewing distance, and *l* = 38 cm. We generated 20 simulated responses (with Poisson noise) for each velocity pairing and computed the depth-sign discrimination index (DSDI, described below) from the responses.

To examine whether depth information can be extracted from model neurons with speed tuning shifted toward head coordinates, we trained a linear decoder to estimate depth *d* from the responses ***r*** of the simulated population:

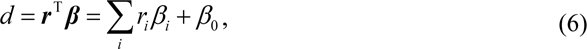

where the weights ***β*** were given by the solution for ordinary least squares:

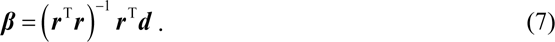

The stimulus parameters used for these simulations were the same as those described in the simulation section of Kim et al. (2017). We generated responses for 500 simulated trials for each stimulus condition to train the decoder, and used responses from another 500 simulated trials as a test set to evaluate decoding performance.

### Depth-sign discrimination index

For both simulated and real neurons, we quantified their depth-sign selectivity using a depth-sign discrimination index, or DSDI (Nadler et al., 2008):

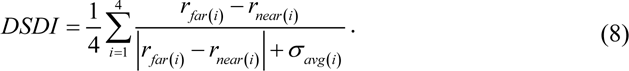

Here, the difference in average firing rate between each pair of far and near depths was calculated relative to the standard deviation of the response to that pair. DSDI was then obtained by averaging across all four pairs of depths, resulting in DSDI∈[−1,1]. Negative DSDI values indicate near-preferring neurons, and positive DSDIs indicate far-preferring neurons. Statistical significance of DSDI values was computed by a permutation test in which the stimulus labels (near or far) for each depth magnitude were shuffled 1000 times. Confidence intervals for DSDI values were obtained by bootstrapping, based on 1000 iterations of resampling with replacement.

### Description of models

We developed a series of models that could be directly fit to the spike trains of each recorded MT neuron. We considered three different extra-retinal components that could modulate the neural response to retinal motion: gain (or multiplicative) modulation (GM), offset (or additive) modulation (OM), and head-centered tuning (HT). Because these three components can operate independently, we are able to examine all possible combinations of them and conduct a factorial model comparison. For all models described in this section, the speed and direction tuning function *f* (·) was the same as in Equation 3.

### Control (Ctrl.) model

As a control, we included a model in which the response is only driven by retinal image velocity:

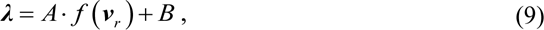

where ***λ*** ∈ ℝ^*N* ×*T*^ represents the firing rates over stimulus duration *T* for *N* trials, *A* is a scaling factor on response amplitude, and *B* is the spontaneous firing rate. The dimensions of the input retinal velocity ***v****_r_* (and eye velocity ***v****_e_*) variable are *N×T×*2. This control model has 7 parameters in total.

### Gain Modulation (GM) model

We constructed a gain modulation model in which the response to retinal velocity is multiplicatively modulated by a function of eye velocity (Figure 8B):

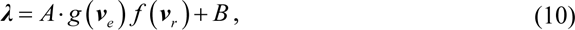

Based on our observations over the range where we have data, the effect of eye velocity is generally monotonic; therefore, we modeled *g* (·) the gain that controls the slope as a sigmoid function with one free parameter *α* that controls the slope:

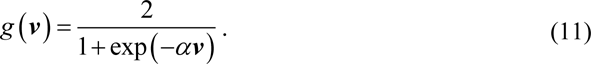

The output of this function is in the range of (0, 2). When the input eye velocity is zero, *g* (0) = 1, and the neural response is therefore determined solely by retinal velocity tuning. Note that this gain function only requires one extra parameter, yielding 8 free parameters in total.

### Offset Modulation (OM) model

We developed another model in which eye velocity has an additive effect on the response to retinal velocity (Figure 8C):

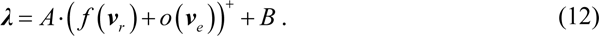

where *o* (·) is an additive offset term and (·)^+^ denotes a rectifier that prevents negative firing rates. Similar to the multiplicative gain function, *g* (·), we modeled the additive effect of eye velocity as a sigmoid with one free parameter *β* :

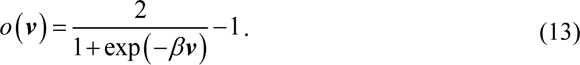

This offset term lies within the range of (-1, 1), thus allowing both suppression and facilitation. This model also has 8 free parameters.

### Head-centered Tuning (HT) model

To test whether MT responses could be explained by velocity tuning in head coordinates, as compared to the GM and OM models, we fit data with the same model used in the simulations described above (Equation 2):

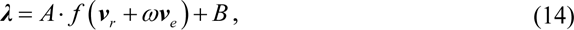

where *ω* is the weight on eye velocity that shifts tuning toward head coordinates, as described in Equations 2 and 4. Like the GM and OM models, this HT model has 8 free parameters.

### Full model and two-component models

To further investigate the role of each model component in accounting for MT responses, we integrated the multiplicative gain *g* (·) term, the additive offset *o* (·) modulation, and the weighting toward head coordinates, *ω*, into one comprehensive model with 10 parameters:

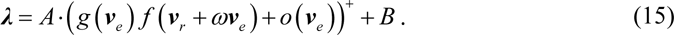

In addition, we can remove each of the components, one at a time, from the full model to create two-component models with 9 free parameters. We refer to these two-component models as –GM, –OM, and –HT, where for example the -GM model is full model without the GM component, such that it contains the OM and HT components.

### Model fitting and comparison

For each neuron, we used spike trains from the main depth-tuning experiments (RM and MP conditions, or RM and DP conditions), as well as from the speed and direction tuning protocols, to fit the above models in two consecutive steps. In the first step, the retinal velocity tuning function (Equation 3) was fit to spike counts from the speed and direction tuning curves. We used bootstrapping (with 1000 iterations of resampling with replacement) to estimate the confidence intervals of each parameter in the retinal velocity tuning function. In the second step, models described in the previous section were fit to spike trains from the main experiments at 1 kHz resolution. 68% and 95% confidence intervals of each tuning parameter estimated in the first step were used here as plausible bounds and hard bounds to constrain the parameters of Equation 3. This fitting procedure acknowledges that there may be overall response differences between direction and speed tuning measurements and the main depth-tuning experiment due to stimulus differences or non-stationary neural responses, while constraining the model fits based on information about tuning curve shapes and stimulus preferences from the independent measurements. We used the MATLAB (Mathworks, MA) function *fmincon* in the first step of the fitting procedure, and Bayesian adaptive direct search, or BADS (Acerbi & Ma, 2017), in the second step. BADS was used to find the global maximum of the log-likelihood log ℒ(**θ**) over *N* trials and stimulus duration *T*:

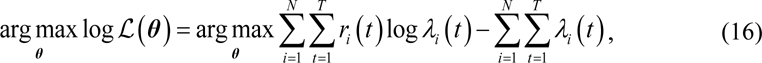

where ***θ*** represents the free parameters in the model being fit, *r_i_* (*t*) represents the actual spikes (0 or 1) at time *t* on the *i*-th trial, and *λ_i_*(*t*) represents the firing rate predicted by the model.

To assess model performance in terms of fitting raw spike trains at 1 kHz resolution, we computed the Bayesian information criterion, or BIC (Schwarz, 1978), for each model:

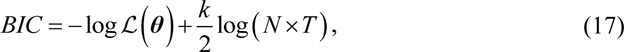

where *k* is the number of free parameters and *N×T* is the number of data points to be fit. To quantify the relative contribution and uniqueness of each model component, we also examined the change of BIC caused by adding or removing each of the three model components (GM, OM, HT). By comparing the GM, OM, and HT models to the control model, we were able to examine how much each component contributes to the fit. By removing one component from the full model (-GM, -OM, -HT models), we were able to measure the uniqueness of the contribution of each component. To assess the absolute goodness of fit of each model, we visually compared the peristimulus time histograms (PSTHs) of neural responses with the corresponding fit of each model. PSTHs were obtained by a moving Gaussian window (*σ* = 33 ms) on the raw spike data and then averaged across repetitions.

Critically, the models were always fit to the spike trains of MT neurons, and we then examined how well the models predicted the depth tuning curves of the MT neurons. Model performance on predicting depth tuning curves was measured by computing the absolute error of the DSDI prediction. Because we only modeled the mean firing rate of each neuron, our models do not directly capture the response variability required for the DSDI calculation (Equation 8). To compute DSDI for each model, we first injected Poisson noise into the model estimates of instantaneous firing rates for each trial, and we then used the simulated Poisson responses to calculate the DSDI for each model from Equation 8. The uncertainty of DSDI predictions was estimated by bootstrapping (1000 resamples with replacement). Since the DSDI predictions were generated solely from the fits to retinal and eye velocities in each trial, the models do not have direct information about depth from MP.

## Results

We analyzed and modeled data from a series of previous experiments on the neural representation of depth from MP in area MT, with the goal of understanding whether the joint neural tuning for retinal and eye velocities is more consistent with gain modulation or head-centered tuning. We examined data from three experimental conditions. In the motion parallax (MP) condition, the animal was laterally translated by a motion platform and counter-rotated their eyes to maintain fixation at a world-fixed target (Figure 1A; Supplementary Figure 1A). A patch of random dots was presented monocularly over the receptive field of an isolated MT neuron, with its motion simulating one of nine depths. In the retinal motion (RM) condition, the animal remained stationary and fixated a static target, while a patch of random dots reproduced the retinal motion that was experienced in the MP condition (Supplementary Figure 1B). Since the random-dot stimulus was generated to be depth-sign ambiguous (Nadler et al., 2008, see Methods), neural responses are not expected to have depth-sign selectivity in the RM condition. In the dynamic perspective (DP) condition, the animal’s posture and the random-dot patch stimulus were the same as in the RM condition, except that a full-field 3D cloud of random-dots was presented in the background (Supplementary Figure 1C). The image motion of these background dots provided an optic flow field that simulated eye rotation (Kim et al., 2015).

We start by exploring the joint tuning of MT neurons for retinal and eye velocities, followed by simulations which show that depth selectivity could emerge from velocity tuning in head coordinates. Next, we fit computational models to MT responses to assess the mechanisms that might account for depth-sign selectivity in both the MP and DP conditions.

### Joint tuning for retinal and eye velocities

As shown by Nadler et al. (2009), the combination of smooth eye movement command signals and responses to retinal motion is sufficient to generate depth-sign selectivity from motion parallax in MT neurons. To better understand exactly how these two signals interact, we visualized how depth from MP maps onto a 2D space having dimensions of retinal velocity and eye velocity. According to the motion-pursuit law (Nawrot & Stroyan, 2009), when the observer is translating laterally and fixating a world-fixed target by compensatory pursuit (Figure 1A), depth from motion parallax is approximately proportional to the ratio of retinal velocity and eye velocity (Equation 5). Hence, depth is defined by the slope of a line that passes through the origin in the 2D velocity plane (Figure 1B).

In the MP condition, retinal and eye velocities co-vary in a quasi-sinusoidal fashion (Figure1 and D; Supplementary Figure 2A-D). Therefore, the idealized trajectory of each trial forms a line (iso-depth line, Figure 1B) whose slope describes the simulated depth of the patch in this trial. Following this logic, we mapped the temporal response in each trial (Supplementary Figure 2E and F) onto the 2D velocity space. Joint tuning for retinal and eye velocities was then estimated by mapping the responses for all trials onto this velocity plane (Supplementary Figure 2G). Note that the eye velocity (Supplementary Figure 2G, x-axis) for the RM and DP conditions represents the eye velocity that was visually simulated while the real eye was stationary. It is also worth noting that the magnitude of MP-based depth scales up rapidly and eventually becomes undefined as the iso-depth line approaches vertical (Equation 5). Since the range of simulated depths in the experiments was limited to the range from -2° to +2° of equivalent disparity, the available data only sample a portion of this joint velocity tuning space. This results in the “bow-tie” shape of the data (Supplementary Figure 2G; Figure 3).

In general, we hypothesize three types of interaction between retinal and eye velocities (Figure 2): partial shifts in tuning curve (Figure 2A), multiplicative gain modulation (Figure 2B), and additive offset modulation (Figure 2C). When eye velocity causes a shift in the retinal tuning curve (Figure 2A), the joint tuning will exhibit a negative diagonal structure (Figure 2D). On the other hand, both multiplicative and additive modulation would expect the response to be suppressed when the eye moves in a particular direction and be facilitated when eye movement is in the opposite direction (Figure 2E and F).

Figure 3A-C shows data from the most typical type of neuron that we found in MT. In the RM condition, since visual cues associated with simulated eye rotation are very weak when the stimulus is small (Kim et al., 2015), the firing rate primarily varies with retinal velocity but not with (simulated) eye velocity (Figure 3A). The corresponding depth tuning curve for the RM condition is thus approximately symmetric around zero-depth (Figure 3C, solid squares). In the MP condition, the response to retinal image motion was facilitated by positive eye velocities and suppressed by negative eye velocities (Figure 3B). In line with previous findings, this pattern is well-described by a multiplicative gain modulation of eye movement on the responses to retinal motion (Kim et al., 2017), such that the joint tuning profile is multiplicatively separable (Figure 2E). This gain modulation leads to a highly asymmetric depth tuning curve (Figure 3C, open circles), yielding a near preference.

We found that other neurons, which prefer very slow retinal velocities and are substantially less common, show a strikingly different pattern of joint tuning. These slow-speed preferring cells exhibit RM responses that peak near zero retinal speed (Figure 3D and G), producing symmetric depth tuning curves with a peak at zero-depth (Figure 3F and I, solid squares). In the MP condition, the retinal velocity tuning of these neurons is neither uniformly suppressed nor facilitated by eye velocity, as expected from gain modulation. Instead, the peak of the retinal velocity tuning shifts systematically with eye velocity, causing a diagonal structure in the joint tuning profile (Figure 3E and H). The diagonal structure, which is a clear deviation from multiplicative separability, produces a preference for far depths (Figure 3F and I, open circles).

This new observation of diagonal structure in the joint tuning suggests an alternative explanation for MP-based depth tuning: some MT neurons might be tuned to velocity in head coordinates (the sum of retinal and eye velocities), instead of being gain-modulated by eye velocity. To better understand the link between this non-separable joint velocity tuning and head-centered velocity coding, consider a scenario in which a stationary observer is pursuing a moving target by eye rotation, while another object is moving in the world (Figure 4A). In this case, the retinal velocity of a moving object reflects both the object motion relative to the world and the image motion induced by the pursuit eye movement. Object velocity in head-centered coordinates can be obtained by simply adding the eye velocity to the retinal velocity. As a result, each negative diagonal line in the retinal velocity vs. eye velocity plane corresponds to a given head-centered speed (Figure 4B). Therefore, a joint tuning velocity tuning profile that follows a negative diagonal (e.g., Figure 3E, H) implies velocity tuning in head coordinates.

### Emergent selectivity for depth from head-centered speed tuning

To examine how depth selectivity could emerge from neurons tuned to motion in head coordinates, we simulated neurons with log-Gaussian speed tuning in a space ranging from retinal to head coordinates, and we calculated depth tuning curves from these model neuron responses.

For simulated neurons that prefer fast speeds (Figure 4C-F), the joint velocity tuning profile in the MP condition produces a broad diagonal band of response. Over the range of retinal and eye velocities sampled in the experiments analyzed here (space between white lines in Figure 4C-E), the predicted result resembles data observed for many of the recorded MT cells (compare Figure 4D, E with Figure 3B), and the corresponding depth tuning curve has a near preference (compare Figure 4F with Figure 3C). Notably, both low and high weights on eye velocity (*ω* = 0.25 and 0.75, respectively) can produce depth selectivity (Figure 4F, blue and orange dashed lines). Consequently, model neurons with tuning shifted toward head coordinates and fast speed preferences produce tuning patterns that are difficult to distinguish from the multiplicatively separable profiles expected from a gain modulation mechanism, given the limited range of the data available.

For model neurons with slow speed preferences (Figure 4K-N), the negative diagonal structure in the joint velocity tuning profile is evident within the range of the measured data (e.g., Figure 4L). This structure produces depth tuning with a far preference (Figure 4N, dash lines), even for relatively small shifts toward head coordinates (low weights on eye velocity). When model neurons have an intermediate speed preference (Figure 4G-J), the preferred depth transitions from far to near as the weight on eye velocity increases (Figure 4J). These results indicate that velocity tuning that is even partially transformed toward head coordinates can predict a wide variety of depth tuning patterns from MP, qualitatively similar to those observed in area MT (Figure 3).

To examine the quantitative relationships between shifts toward head coordinates and depth-sign selectivity, we sampled the parameters in our model within a biologically plausible range, and we systematically varied the weight on eye velocity to control the degree of shift in reference frame toward head coordinates. We generated Poisson spikes from the simulated neurons, and used them to compute a depth-sign discrimination index (DSDI, see Materials and Methods for details) as a quantitative measurement of depth selectivity, as used previously in empirical studies (e.g., Nadler et al., 2008; Nadler et al., 2009). Surprisingly, significant depth selectivity (*p* < .05, permutation test; Figure 5A) starts emerging when the weight on eye velocity is as low as 0.01 and saturates around 0.2, suggesting that robust depth tuning can arise from even a modest shift toward head-centered speed tuning.

Our simulation also reveals a striking relationship between preferred speed and depth-sign preference, as suggested by the example units of Figure 4. To further investigate this relationship, we simulated a population of neurons with preferred speeds that are equally spaced on a logarithmic scale from 0.1 to 30 °/s. We find a strong relationship between the preferred speed in head coordinates and preferred depth (Figure 5B, such that neurons with slow speed preferences tend to prefer far depths (Spearman’s *r* = -0.748, *p* = 8.47×10^-91^). A very similar pattern was reported in the empirical data of Nadler et al. (2008) (Spearman’s *r* = -0.561, *p* = 3.06×10^-13^; Figure 5C), which previously had no clear explanation. To examine whether the relationship between preferred speed and depth-sign preference is also predicted by multiplicative gain modulation or additive offset modulation, we simulated these mechanisms based on Equations 10 and 1 using the same range of parameters. We find no correlations between preferred speed and DSDI in either cases (Spearman’s *r* = -0.035, *p* = .436 for gain modulation; Spearman’s *r* = 0.076, *p* = .089 for offset modulation; Supplementary Figure 3). We show that this pattern of DSDIs only emerges naturally from a representation that is partially transformed toward head coordinates, which suggests that this mechanism might underlie the depth-sign selectivity previously described.

We further explored whether a neural population with such head-centered speed tuning could be used to accurately estimate depth from MP. Poisson spikes were generated from neurons with velocity tuning that was shifted toward head coordinates, with the preferred speeds and weights on eye velocity sampled from a logarithmic space having ranges of [0.1, 30] and [0.001, 1], respectively. Other parameters including tuning width, response amplitude, and baseline firing rate were randomly sampled from ranges that are constrained by empirical data (see Materials and Methods for details). We trained a simple linear decoder on the population response for 500 simulated trials to recover the magnitude and sign of depth. The performance of the decoder was then evaluated by predicting depth from the responses in another 500 test trials. The linear decoder successfully estimated depth in the MP condition, while it performed very poorly in the RM condition (Figure 5D). The decoder also failed when model responses for the MP condition were shuffled across combinations of eye velocity and retinal velocity (Figure 5D, purple dashed line). These results demonstrate that a population of neurons with tuning that is shifted toward head-centered velocity can reliably represent depth from MP.

### Modeling neural responses with head-centered tuning and gain modulations

While head-centered speed tuning could account for the empirical findings of depth selectivity, it remains unclear whether the data generally support this head-centered tuning hypothesis over the previous theory of gain modulation (Kim et al., 2017). To quantitatively compare these hypotheses, we evaluated models that incorporated three components that describe types of interactions between eye velocity and retinal velocity: a shift toward head-centered tuning (HT), multiplicative gain modulation (GM), and additive offset modulation (OM) (Supplementary Figure 4). As described above, the HT model incorporates a weighted sum of retinal and eye velocities as the input to the speed and direction tuning function, *f* (·) (Supplementary Figure 4A). By contrast, the tuning function, *f* (·), in both GM and OM models only takes retinal velocity as input, while its output is modulated by eye velocity in either a multiplicative or additive manner (Supplementary Figure 4B and C). These three components are not mutually exclusive and were also combined together in different ways. We therefore considered all eight combinations of these model components, including a control model (Ctrl.) in which there is no influence of eye velocity, three models with only one interaction component (HT, GM, and OM), a full model with all three components (Full), and three models with one of the components removed (–HT, –GM, and –OM). The velocity tuning function in each model was first fit to speed and direction tuning measurements from each neuron (e.g., Figure 6A) to determine parameter bounds, and then each model was fit to spike trains from the main experimental conditions (RM and MP conditions). We then used the estimated parameters of each model to predict depth tuning curves and DSDI values. Our examination of all combinations of model components allows us to assess the commonalities and uniqueness of each hypothesized mechanism.

**Figure 6.**
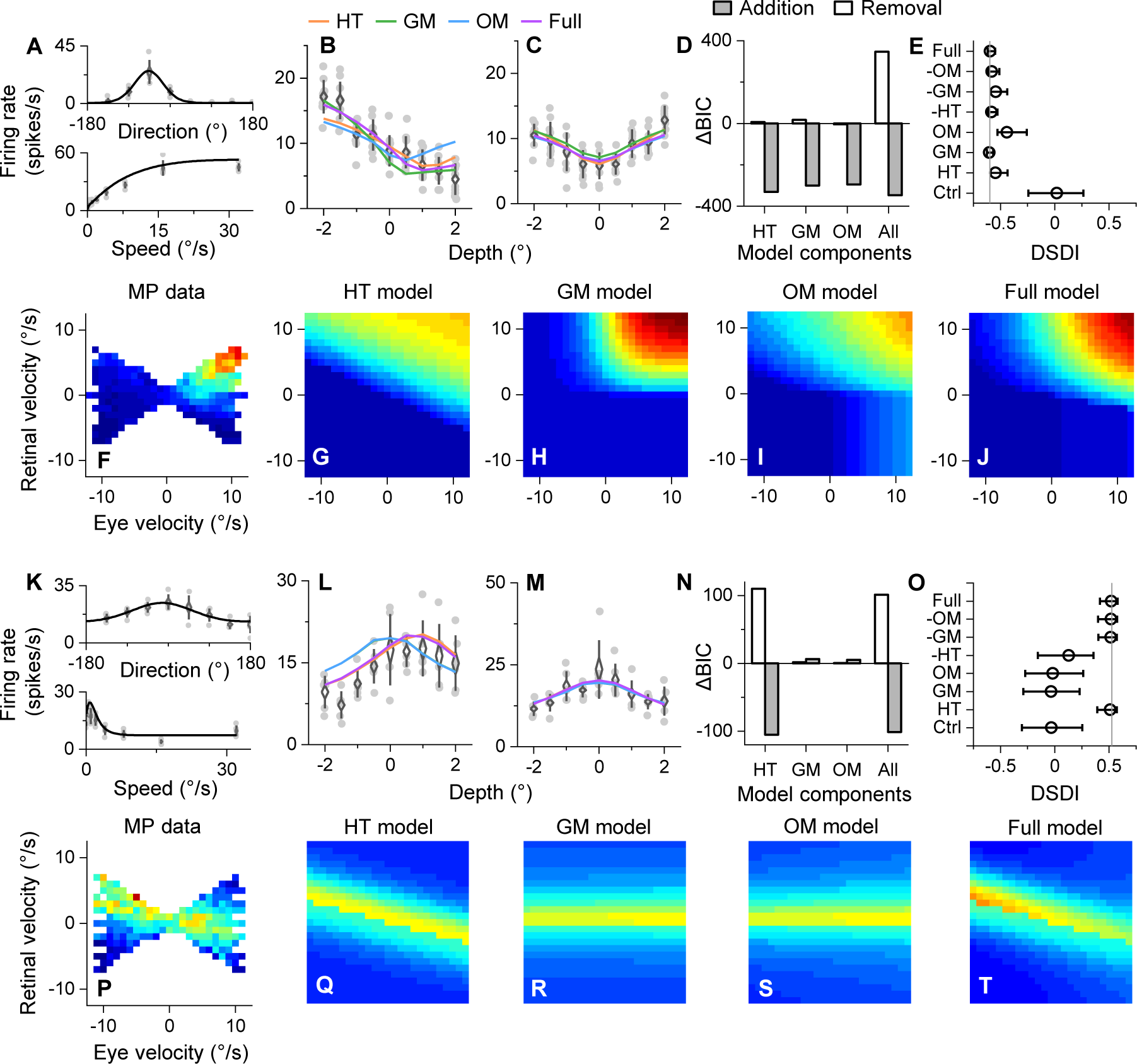
Examples of model fitting results. **A-J**, Model fitting results for a neuron that prefers fast speed (the same neuron as shown in Fig. 3A-C). **A**, Retinal direction (upper panel) and speed (lower panel) tuning functions were fit to data from the tuning measurements. Gray dots indicate mean firing rates of each trial, black curve shows the best fit of the model. Error bars show 1 SD around the mean of the data. **B** and **C**, Depth tuning in both MP (B) and RM (C) conditions and the corresponding model predictions (orange curve, HT model; green curve, GM model; blue curve, OM model; purple curve, Full model). Gray dots indicate the mean firing rates of each trial and open circles represent the averaged firing rates across repetitions. Error bars represent 1 SD around the mean. **D**, Change of BIC after adding each component to the control model (gray bars) or removing them from the full model (white bars). Negative change of BIC indicates better fitting. **E**, DSDI predicted by each model. Vertical line shows the actual DSDI in the MP condition. Error bars show 95% confidence intervals estimated by 1000 times bootstrapping. **F**, Joint tuning in the MP condition. **G-J**, model predictions of the joint tuning pattern in the MP condition (**G**, HT model; **H**, GM model; **I**, OM model; **J**, full model). **K-T**, Model fitting results for a neuron that prefers slow speed and far depth (the same neuron as shown in Fig. 3D-F).

Figure 6A-J shows the fitting results for a typical near-preferring neuron with a relatively high-speed preference. We found that all models fit to spike trains captured the temporal structure of response histograms reasonably well (Supplementary Figure 5A), and all models predicted the depth tuning reasonably well (Figure 6B and C). To quantify the uniqueness of each model component, we computed the change of the Bayesian Information Criterion (ΔBIC) in two different ways: 1) by adding each model component to the control model (gray bars, Figure 6D), and 2) by removing each component from the full model (white bars, Figure 6D). For this type of neuron, adding any of the three components to the Ctrl model greatly improved the goodness-of-fit of spike trains, while removing any individual component showed little reduction in BIC (Figure 6D). This indicates that each of the three model components (HT, GM, and OM) is largely capable of capturing the data for this neuron. Analysis of the model predictions of DSDI reveals a similar result: all models except the control model predicted the neuron’s depth-sign selectivity pretty accurately (Figure 6E). One can appreciate why all of the models fit the data well by examining the raw data and fits in the joint velocity space (Figure 6F-J). Because the speed preference is relatively high, the empirical data are missing in the region of the joint velocity space (high retinal speeds and low eye speeds) that would best distinguish among models, especially HT (Figure 6G) versus GM (Figure 6H). Thus, HT, GM, and OM models are equally likely to explain this type of joint tuning pattern given the limitations of the empirical data.

In contrast, the model fits to responses of neurons with clear diagonal structure in their joint velocity profile show a different pattern of results (Figure 6K-T; Supplementary Figure 5G-L). For these cells, the HT model clearly outperformed the GM and OM models in both fitting spike trains and in predicting depth tuning curves (Figure 6L-O; Supplementary Figure 5G). In addition, the HT model uniquely accounted for the diagonal structure in the joint velocity tuning (Figure 6P-T), such that removing the HT component resulted in a large increase in BIC and DSDI prediction error (Figure 6N and O). This result further supports our hypothesis that some neurons have velocity tuning that is shifted toward head-centered coordinates, and that this shift produces a far preference for depth.

We compared the models at the population level based on both ΔBIC values and DSDI prediction errors. For the component addition manipulation, a negative ΔBIC indicates that a better fit is obtained by adding the model component to the control model. In general, there was a negative ΔBIC when a component was added to the control model, with significant variation across components (*χ*^2^(3) = 307.02, *p* = 3.01×10^-66^, Kendall’s W = 0.463, Friedman’s test; Figure 7A). Across the population, the ΔBIC value obtained by adding either the HT or OM component is significantly more negative than that obtained by adding the GM component (*Z* = -9.306, *p* = 1.33×10^-20^ for HT-GM; *Z* = -9.157, *p* = 5.34×10^-20^ for OM-GM; Wilcoxon signed-rank test; Bonferroni corrected *α* = .0167; Figure 7A), and no significance was found in ΔBIC between the HT and OM models (*Z* = 1.850, *p* = .0643, Wilcoxon signed-rank test). The relative contribution of each model component to ΔBIC for individual neurons is visualized in Supplementary Figure 6A, where each vertex represents a dominant contribution by one component. Most neurons show substantial contributions of the HT and OM components, while fewer cells have a strong contribution of the GM component. As a result, the largest density of neurons lies in the lower-middle region of the triangle.

**Figure 7.**
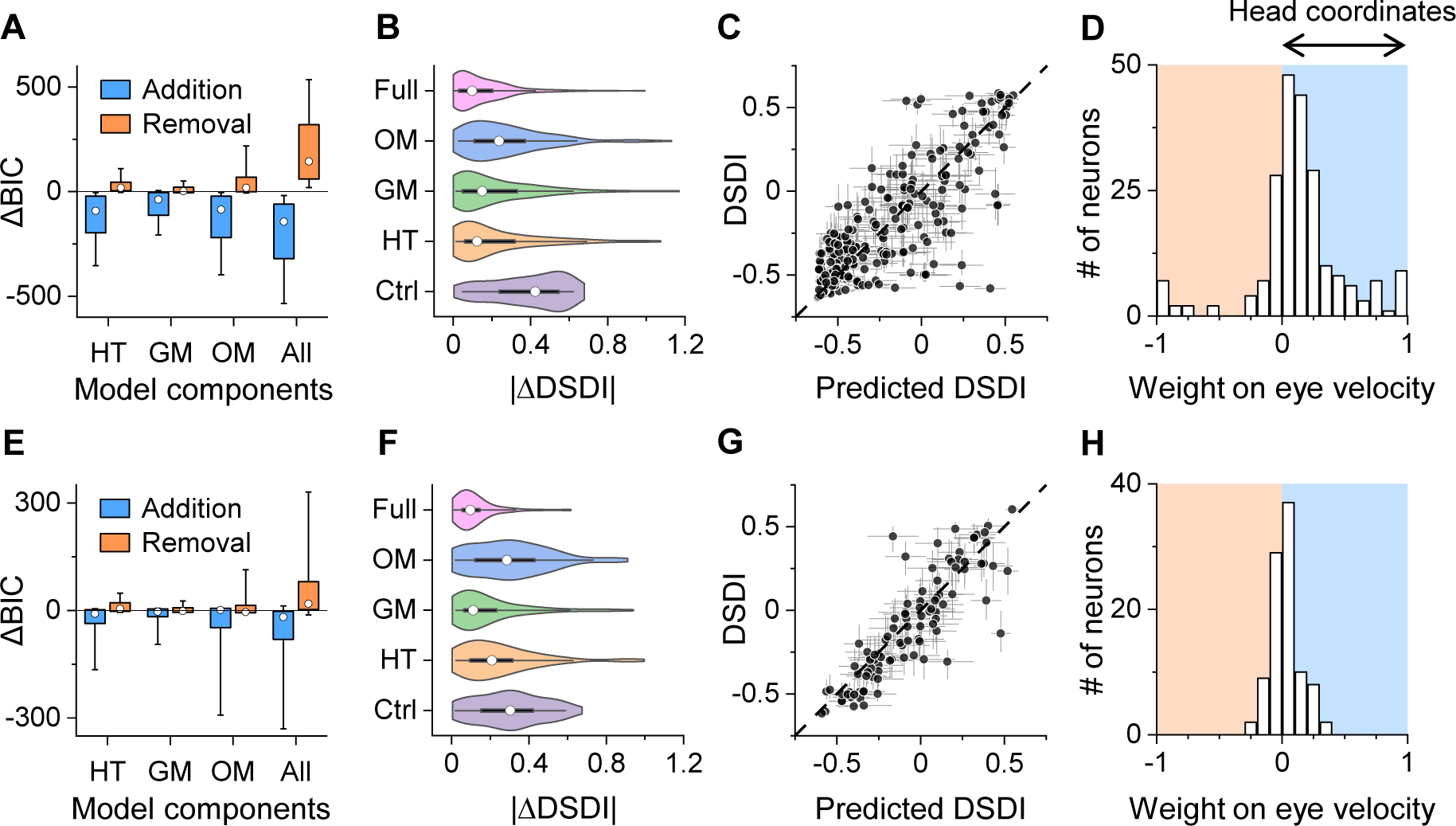
Summary of model fitting results in MP and DP conditions. **A-D**, Summary of results in the MP condition. **A**, Change of BIC by addition and removal for all neurons. Open circles indicate the median, boxes represent interquartile range and error bars indicate 90% CI. **B**, Distribution of |ΔDSDI|. Open circle represents the median, solid black box indicates the interquartile range, and whisker shows the 95% CI. **C**, Scatter plot shows the relationship between DSDI predicted by the full model (x-axis) and the actual DSDI (y-axis). Each dot indicates one neuron; error bars represent 68% CI. **D**, Distribution of the weight on eye velocity in the full model. Positive weights indicate representation in head coordinates. **E-H**, Summary of fitting results in the DP condition.

**Figure 8.**
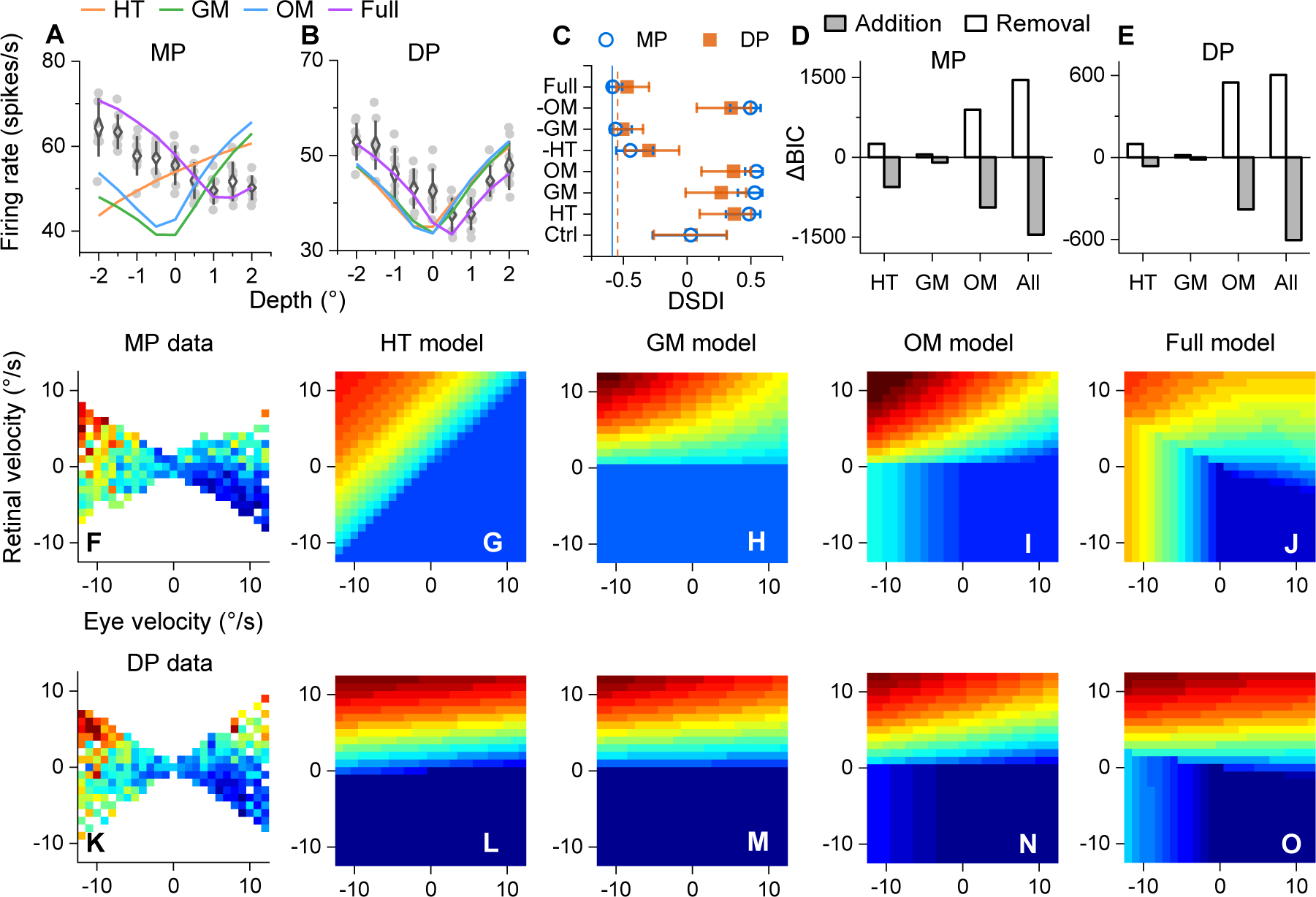
Example joint tuning and model fitting in the DP condition. **A** and **B**, depth tuning curves in MP and DP conditions, respectively. Error bars indicate 1 SD around the mean. **C**, DSDI predictions in both MP (open blue circles) and DP (solid orange squares) conditions. **D** and **E**, Change of BIC by addition (gray bars) and removal (white bars) in MP and DP conditions, respectively. **F**-**J**, Joint tuning in the MP condition (**F**) and the prediction by models (**G**, HT model; **H**, GM model; **I**, OM model; **J**, full model). **K**-**O**, Joint tuning in the DP condition (**K**) and corresponding model predictions (**L-O**).

For the component removal manipulation, a positive ΔBIC indicates that the goodness-of-fit is impaired by removing the component, indicating a unique contribution of that component to the fit. Similar to component addition, there was a significant difference in ΔBIC depending on which model component was removed from the Full model (*χ*^2^(3) = 383.37, *p* = 8.84×10^-83^, Kendall’s W = 0.578, Friedman’s test; Figure 7A). We found a significantly greater ΔBIC when removing the HT component as compared to GM (*Z* = 7.259, *p* = 3.91×10^-13^, Wilcoxon signed-rank test; Bonferroni corrected *α* = .0167), and also for OM as compared to GM (*Z* = 8.066, *p* = 7.29×10^-16^, Wilcoxon signed-rank test). In addition, ΔBIC for removing the OM component is slightly greater than that for HT (*Z* = 2.738, *p* = .0062, Wilcoxon signed-rank test). Together, the ΔBIC data of Figure 7A and Supplementary Figure 6A indicate comparable contributions of HT and OM components to fitting MT responses, with a somewhat weaker contribution from the GM component.

To examine the contribution of model components to predicting depth-sign selectivity, we quantified their performance as the absolute error between the predicted and measured DSDI values, which we denote |ΔDSDI|. We find that median |ΔDSDI| values differ significantly across models (*χ*^2^(4) = 112.46, *p* = 2.18×10^-23^, Kendall’s W = 0.127, Friedman’s test; Figure 7B). We conducted a Wilcoxon signed-rank test for each pair of models and applied a Bonferroni correction, resulting in a significance level *α* = .005 (Table 1). We find that all models predicted depth tuning significantly better than the control mode, and that the full model significantly outperformed all single-component models. In addition, we find that both HT and GM models predict depth tuning better than the OM model, while there is no significant difference between HT and GM models (Table 1; Figure 7B). The relative proportions of |ΔDSDI| from each model are shown in Supplementary Figure 6B. Note that there is a lower density of cells around the vertices of GM and HT components, as compared to OM, indicating that GM and HT components predicted depth selectivity more accurately for most neurons (Supplementary Figure 6B). Finally, the Full model performed well in predicting the depth selectivity for most MT neurons (Spearman’s *r* = 0.803, *p* = 3.87×10^-51^; Figure 7C), which is especially notable given that the model was fit to the spike trains of neurons and was not optimized to predict depth-sign selectivity per se.

**Table 1.** Results of Wilcoxon signed-rank tests comparing |ΔDSDI| values for pairs of models in the MP condition. The significance level after Bonferroni correction is 0.005.

| MODEL PAIRS | Z-VALUE | P-VALUE |
| --- | --- | --- |
| HT – CTRL | -6.948 | $3.72 \times 10^{-12}$ |
| GM – CTRL | -6.861 | $6.82 \times 10^{-12}$ |
| OM – CTRL | -6.145 | $8.01 \times 10^{-10}$ |
| FULL – CTRL | -10.013 | $1.34 \times 10^{-23}$ |
| HT – FULL | 4.249 | $2.15 \times 10^{-5}$ |
| GM – FULL | 4.126 | $3.69 \times 10^{-5}$ |
| OM – FULL | 7.254 | $4.04 \times 10^{-13}$ |
| HT – GM | 0.988 | .323 |
| HT – OM | -3.701 | .0002 |
| GM – OM | -4.694 | $2.69 \times 10^{-6}$ |

During the fitting of the HT component, we constrained the weight on eye velocity to the range from -1 to 1. As shown in Equation 4, this parameter reflects the shift of tuning toward head coordinates when it has a positive value. Indeed, we found that most of the estimated weights are biased toward positive values (median = 0.122, *Z* = 7.830, *p* = 2.44×10^-15^, one-tailed Wilcoxon signed-rank test; Figure 7D), indicating that this weight term is generally meaningful for representing head-centered velocity, rather than overfitting an arbitrary data pattern.

### Modulation by simulated eye velocity from large-field background motion

In this section, we examine whether the mechanisms of interaction between retinal velocity and eye velocity are similar when eye movements are simulated by large-field visual background motion while the eye remains stationary. Specifically, we analyzed data from the dynamic perspective (DP) condition (Supplementary Figure 1C) in the same way as we did for the MP condition.

Figure 8 shows data from two example neurons and the corresponding model fits. The first example neuron exhibits a clear additive contribution of eye velocity (evidenced as a vertical stripe-like pattern on the left side of the joint velocity tuning profiles, Figure 8F, K), which is captured by models with an OM component. As a consequence, the OM component contributes most to ΔBIC values in both the MP and DP conditions (Figure 8D and E). However, the OM component alone could not explain the observed depth tuning curves (Figure 8A and B). Only models with both OM and one other component (either HT or GM) were able to correctly predict the sign of depth-sign selectivity (Full, –HT, and –GM models in Figure 8C). On the other hand, the second example neuron in Supplementary Figure 7 shows a more gain-like pattern of modulation in both MP and DP conditions, with no clear signatures that would suggest the necessity of either an OM or HT component (Supplementary Figure 7). Thus, all three model components fit the data similarly well, and are mostly interchangeable (Supplementary Figure 7A-E). For both example neurons, the joint velocity tuning and depth tuning are quite similar between the MP and DP conditions, consistent with previous findings (Kim et al., 2015). However, we were not able to identify any neurons with a clear negative diagonal structure in the joint velocity tuning profile for the DP condition. This might be due to the fact that fewer neurons were found to prefer far depths and slow speeds in this dataset, or to a difference between real and simulated eye movements (see Discussion).

In general, population results for the DP condition (Figure 7E-H) are similar to those for the MP condition (Figure 7A-D). We found a significant dependence of ΔBIC on which model component was added or removed (*χ*^2^(3) = 40.11, *p* = 1.01×10^-8^, Kendall’s W = 0.138 for addition; *χ*^2^(3) = 46.14, *p* = 5.30×10^-10^, Kendall’s W = 0.159 for removal; Friedman’s test; Figure 7E; Supplementary Figure 6C). ΔBIC values were significantly different between the HT and GM components (*Z* = -4.939, *p* = 7.87×10^-7^ for addition; *Z* = 4.546, *p* = 5.46×10^-6^ for removal; Wilcoxon signed-tank test; Bonferroni corrected *α* = .0167), but not between other pairs of components (*Z* = -1.355, *p* = .176 for additions of HT and OM; *Z* = 1.589, *p* = .112 for removals of HT and OM; *Z* = 1.204, *p* = .229 for additions of GM and OM; *Z* = -0.272, *p* = .786 for removals of GM and OM; Wilcoxon signed-tank test).

Significant differences between models were also found for |ΔDSDI| values in the DP condition (*χ*^2^(4) = 60.08, *p* = 2.79×10^-12^, Kendall’s W = 0.155, Friedman’s test; Figure 7F; Supplementary Figure 6D). Wilcoxon signed-rank tests with Bonferroni correction for multiple comparisons were again performed to compare |ΔDSDI| between each pair of models (Table 2). In contrast to what we found for the MP condition, only the GM and Full models showed significantly better predictions of DSDI than the control model, while the Full model outperformed all other models. Our Full model also performed very well in predicting the depth-sign selectivity of MT neurons in the DP condition (Spearman’s *r* = 0.884, *p* = 4.56×10^-33^; Figure 7G).

**Table 2.** Results of Wilcoxon signed-rank tes

| MODEL PAIRS | Z-VALUE | P-VALUE |
| --- | --- | --- |
| HT – CTRL | -2.747 | .0060 |
| GM – CTRL | -4.313 | **$1.61 \times 10^{-5}$** |
| OM – CTRL | -1.148 | .251 |
| FULL – CTRL | -6.633 | **$3.28 \times 10^{-11}$** |
| HT – FULL | 5.439 | **$5.37 \times 10^{-8}$** |
| GM – FULL | 3.024 | **.00249** |
| OM – FULL | 6.399 | **$1.56 \times 10^{-10}$** |
| HT – GM | 3.891 | **$9.96 \times 10^{-5}$** |
| HT – OM | -2.909 | **.00362** |
| GM – OM | -5.108 | **$3.26 \times 10^{-7}$** |

We also compared the model performance in predicting DSDIs between the MP and DP conditions. The performance of the Full model, based on |ΔDSDI| values, did not differ significantly between these two conditions (*Z* = -0.235, *p* = .815, Wilcoxon rank-sum test). A modestly significant difference between MP and DP conditions was only found for the HT model (*Z* = 2.137, *p* = .033, Wilcoxon rank-sum test), but not for the GM or OM models (*Z* = -0.289, *p* = .773 for GM; *Z* = 1.488, *p* = .137 for OM; Wilcoxon rank-sum test). We also found that the weights on eye velocity in the HT component are closer to zero and less biased toward positive values in the DP condition (median = 0.0083, *Z* = 1.979, *p* = .024, one-tailed Wilcoxon signed-rank test for zero medians; *Z* = 5.360, *p* = 4.17×10^-8^, one-tailed Wilcoxon rank-sum test for smaller median in DP as compare to MP condition; Figure 7H). This indicates that shifts toward head-centered velocity tuning are less prominent in the DP condition.

## Discussion

Our findings show that depth selectivity from motion parallax in area MT can arise from even a modest shift in velocity tuning from retinal to head coordinates. While the joint velocity tuning of many MT neurons is consistent with the previous suggestion of a gain modulation mechanism (Kim et al., 2017), other neurons with slow speed preferences show a clear shift in retinal speed preference with eye velocity that manifests as a diagonal structure. Our simulations reveal that a range of depth tuning properties can be explained by a partial shift of velocity tuning toward head coordinates. We further demonstrate that a population of such neurons can be simply decoded to estimate depth, and that the previously observed relationship between depth-sign selectivity and speed preference (Nadler et al., 2008) emerges naturally from a tuning shift toward head coordinates. Our models performed well in fitting spike trains and predicting depth-sign selectivity of MT neurons, and the HT component uniquely contributed to the fitting of a subset of MT neurons. However, it must be noted that the empirical data are not sufficient to distinguish among models for many MT neurons, due to the limited range of retinal and eye velocities tested. Together, our results open the possibility that depth-sign selectivity in area MT might arise from a mechanism that may be involved in representing object motion in head coordinates during self-motion.

### Relationship to previous studies on depth coding from MP

This study extends previous work on MP-based depth coding in the following ways. First, we analyzed raw spike trains at millisecond resolution, whereas previous analyses were generally performed on mean spike rates over a one- or two-second time period (Kim et al., 2015, 2017; Nadler et al., 2008). This fine-scale analysis allowed us to examine detailed response modulations by different magnitudes of eye velocity, thus providing estimates of joint tuning for retinal velocity and eye velocity. Second, we formalized a gain modulation model based on previous theory, and addressed its strengths and weaknesses in explaining joint velocity tuning patterns. Third, we proposed a new computational model of head-centered speed tuning based on our observations, and compared it with a variety of other model variants in a rigorous manner. Our HT model not only successfully accounts for specific patterns of joint velocity tuning that the GM and OM models fail to explain, but it also naturally predicts a previously unexplained population relationship between preferred speed and depth-sign selectivity across a population of neurons (Nadler et al., 2008).

The HT, GM and OM models performed similarly well in fitting the responses of many MT neurons, suggesting the possibility of multiple mechanisms that may be somewhat redundant. In Kim et al. (2017), a large portion of MT neurons was found to be gain modulated by the direction of eye movement in a multiplicative fashion. However, that study did not explicitly consider other models nor compare models in a systematic fashion. Indeed, for many neurons, our results show that the HT model can produce joint tuning profiles similar to those predicted by gain modulation, especially given the limited range of the experimental data. These results imply that the gain-modulation effects reported previously by Kim et al. (2017) might also arise from a shift in velocity tuning toward head-centered coordinates. In addition, for a subset of MT neurons with slow speed preferences, our findings demonstrate clearly that depth tuning arises from a shift toward head-centered tuning, not gain modulation. However, it remains to be determined whether head-centered tuning also accounts for the selectivity of most MT neurons with higher speed preferences. It is also possible that multiple mechanisms co-exist in MT and produce similar depth-sign selectivity for motion parallax.

### Differences between MP and DP conditions

In this study, we fit the same set of models to neural responses recorded in both the MP (real eye movement) and DP (simulated eye movement) conditions. While the pattern of results across the population is highly similar between these two conditions (Figures 7), we find that weights on eye velocity in the HT model for the DP condition concentrate more around zero and are less biased toward positive values. This implies that a shift toward a head-centered representation of speed may be less prominent in the DP condition. We speculate that this arises from the different levels of ambiguities of extra-retinal signals in the MP and DP conditions. As shown by Nadler et al. (2009), a smooth pursuit command signal, but not a vestibular signal, is used to disambiguate MP-defined depth. When an eye movement signal is the only extra-retinal signal in use, the source of retinal motion is ambiguous: it could be the result of self-motion or object motion in the world. When the scene is interpreted in the latter way, head-centered speed tuning may naturally occur as an effort to infer object motion in the world. However, the large-field visual background motion in the DP condition provides an unambiguous cue to distinguish between these two scenarios. Both translational and rotational optic flow will be produced when the observer is stationary and pursuing a moving target, while only rotational background motion occurs when the observer translates laterally and fixates a world-fixed target. This means that the extra-retinal signal in the DP condition is not ambiguous, and the animal can infer the true geometry (self-motion with eye rotation) without misinterpreting the scene. Thus in the DP condition, head-centered speed representation may not be required by the computation.

### Comparison with previous work on reference frames of speed tuning in MT

Both Chukoskie and Movshon (2009) and Inaba et al. (2011) measured the speed tuning of neurons in MT and MST during fixation and during smooth pursuit eye movements. They reported that MT neurons generally represent motion in retinal coordinates, whereas MST neurons are typically tuned to screen coordinates. However, at least some MT neurons in their data reveal a clear response shift toward screen coordinates, which is equivalent to head coordinates since neither the head nor the screen was moving in their experiment. Our simulations demonstrate that even a slight shift towards head coordinates can produce robust selectivity for depth from MP, and our model-fitting results confirmed that the shifts toward head coordinates are generally quite small. Thus, while the extent of tuning shifts toward head coordinates in our MT data are compatible with previous findings, these small shifts appear to be sufficient to account for the depth-sign selectivity of MT neurons.

### Limitations of the present study

Since the magnitude of depth from MP is unrealistically large when the iso-depth line is close to vertical (Figure 1B), the empirical data that we analyzed has limited samples from the entire joint velocity space. Our model fitting results reveal that differences among the HT, GM and OM models are subtle for many neurons over the range of available data. Therefore, a more extensive sampling of the joint tuning for retinal and eye velocities will be necessary to distinguish these different mechanisms. Experiments that are not constrained by the geometry of MP (Figure 1B) might be useful for measuring the interaction between retinal and eye velocities across much wider ranges. Note that our models were fit to spike trains without accounting for delays or dynamics in the neural responses. We explored modeling the temporal dynamics of neural responses, but found that it made little difference to the fits since the temporal structure of our stimuli is slow (0.5 Hz movement). From a system identification perspective, uncorrelated white noise signals may be better suited for measuring MT responses to a range of retinal and eye velocities (Brostek et al., 2015), as well as for building a model that fully describes the bottom-up mapping from velocity signals to neural responses in MT.

We observed that a small proportion of the recorded neurons (< 5%) showed joint velocity tuning profiles that were dominated by eye velocity, and this could be observed for both DP and MP conditions (Supplementary Figure 8A and F, respectively). While these neurons show clear retinal speed tuning during a fixation task, this retinal velocity contribution to the joint tuning is overwhelmed by the effect of (real or simulated) eye velocity for this small subset of neurons. None of our model components could explain this observed pattern adequately (Supplementary Figure 8B-E and G-J), and we speculate that a normalization mechanism might be able to explain these data, as previous work has shown that normalization can approximate a winner-take-all interaction when inputs are imbalanced (Busse et al., 2009).

We have only explored the neural interactions between retinal velocity and eye velocity in the context of an observer who translates laterally while maintaining fixation at a world-fixed target (Figure 1A). We have suggested that this viewing context could potentially be misinterpreted as a coordinate transformation (CT) context in which a stationary observer tries to infer object motion in the world by compensating for their eye movements (Figure 4A). However, it is unclear how exactly MT neurons jointly encode retinal and eye velocities when directly tested in the CT context. While we showed that some neurons might represent velocity in head coordinates, we have not directly examined the link between head-centered velocity tuning and the CT context that explicitly requires this kind of representation. It would be interesting to see whether the diagonal structure in the joint velocity tuning profile of MT neurons is observed more frequently if an animal is trained to judge the head-centered speed of an object during pursuit eye movements. Comparing the neural interaction of retinal and eye velocities in different task contexts will also provide insights into how much the computations performed by MT neurons are shaped by top-down task-dependent signals.

### Implications for the role of MT in inferring object motion and depth during self-motion

Theories of sensorimotor transformation have shown that coordinate transformation from retinal to head coordinates can be achieved at the *population* level by a group of neurons that are gain modulated by eye position (Pouget & Sejnowski, 1997; Salinas & Abbott, 1995; Salinas & Thier, 2000; Zipser & Andersen, 1988). Our study provides additional evidence that this kind of coordinate transformation not only happens at the population level, but can also occur in *individual* MT neurons to some extent. The diagonal structure of joint velocity tuning that we found in some neurons could be a result of feedback signals from higher-level sensory/motor areas that represent speed in head coordinates (Deneve et al., 2001). It could also emerge from the lateral recurrent connections between neurons in MT, or through the feedback of eye velocity signals that are combined with retinal motion within MT. Both gain modulation and speed tuning that partially shifts toward head coordinates might serve as basis functions that support computations of multiple sensory variables (Deneve et al., 2001; Pouget & Sejnowski, 1997; Salinas & Abbott, 1995). Further studies will be needed to understand the mechanisms that give rise to velocity selectivity in head coordinates, as well as how it shapes the functional contributions of area MT.

Our analyses suggest a shared representation between depth from MP and object motion in the world, indicating that MT is a candidate brain area for studying perceptual inferences that depend on these two variables. It would be interesting to see whether MT responses differ when an animal is trained to perform depth- or motion-related tasks using a common set of stimuli. Neurons might show diverse patterns of joint tuning for the same set of stimuli when the task context changes. If MT neurons indeed convey information about both depth from MP and object motion in the world, another important question is how downstream brain areas selectively read out these variables from the MT population based on task demands.

In summary, this study set out to gain a better understanding of the interactions between retinal velocity and eye velocity that give rise to depth-sign selectivity from motion parallax in MT. By developing computational models that capture the modulation effects of eye velocity, we have demonstrated that MP-based depth-sign selectivity could also emerge from velocity tuning that is at least partially shifted toward head coordinates. These findings highlight the potential role of MT in representing higher-level sensory variables, including depth from MP and object motion in the world.

## Acknowledgments

This work was supported by NIH grant EY013644 (to GCD), and an NEI CORE grant (EY001319).

## Figure legends

**Supplementary Figure 1.**
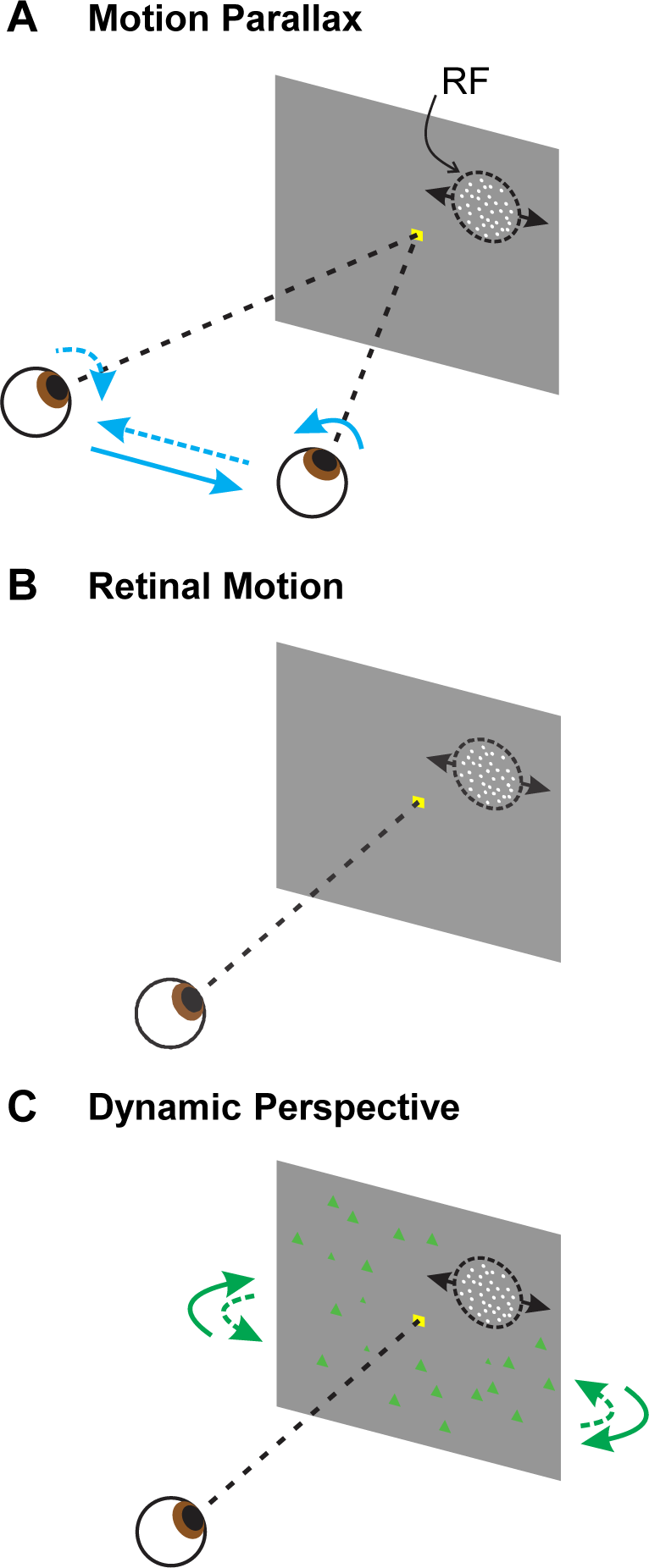
Illustration of experimental conditions. **A**, Illustration of the motion parallax (MP) condition. The animal was laterally translated by a motion platform, while maintaining fixation at a stationary target (yellow square) by making smooth pursuit eye movements. A patch of random dots was presented monocularly and moved over the neuron’s receptive field (RF), simulating the motion parallax of near or far objects. **B**, Retinal motion (RM) condition. The animal stayed stationary and fixated at a static target at the center of the screen. The retinal motion of the patch is a replication of that in the MP condition. **C**, Dynamic perspective (DP) condition. In addition to the RM condition, a large-field 3D background was present, providing visual information about eye movement.

**Supplementary Figure 2.**
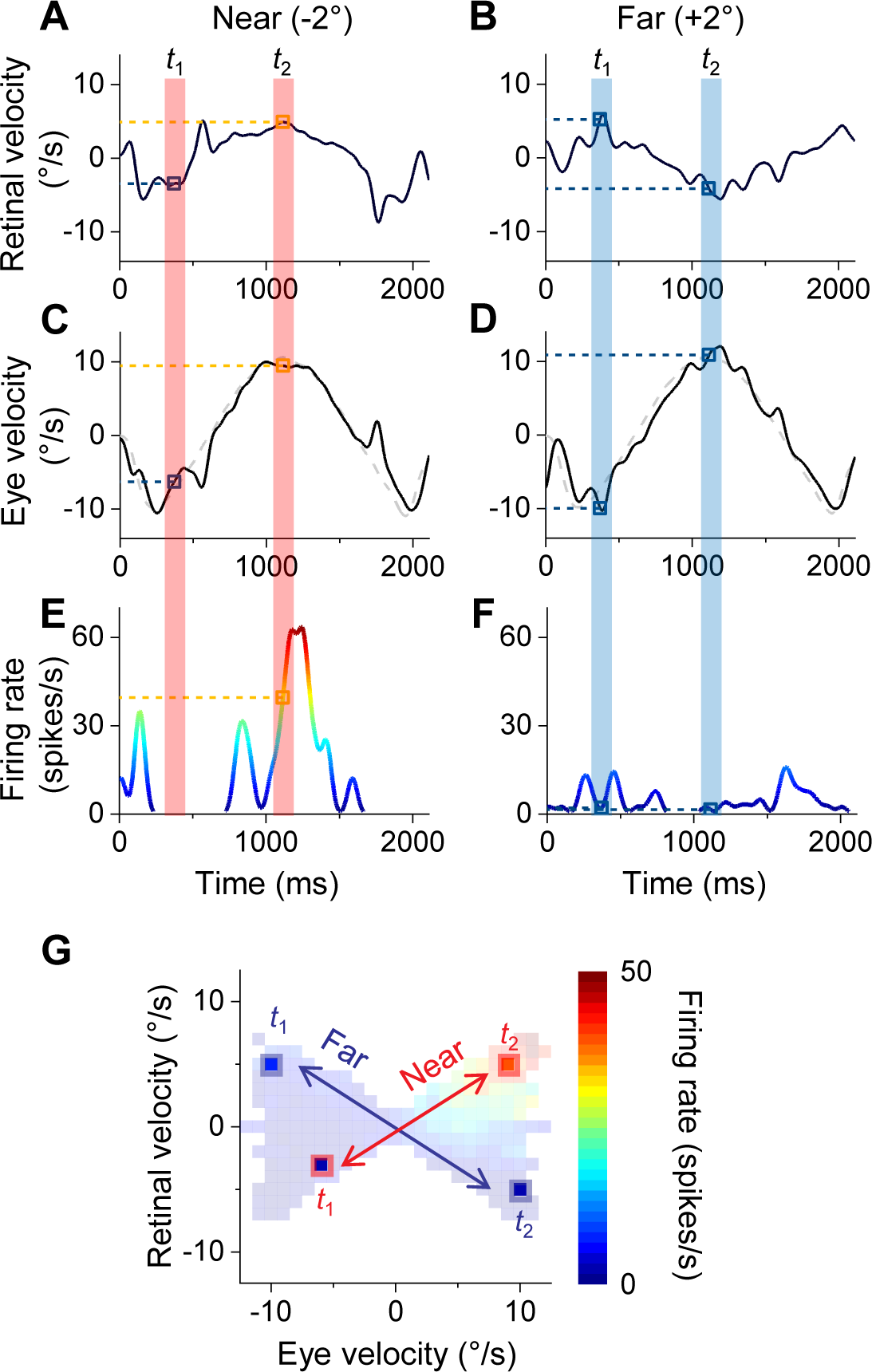
Mapping temporal responses into joint retinal-eye velocities tuning. **A** and **B**, Retinal velocity in near (-2°) and far (+2°) depth, respectively. **C** and **D**, Trajectories of eye velocity (black curve) and target velocity (gray curve) at one example trial for near and far depths. **E** and **F**, PSTHs of example trials in the MP condition for near and far depths. Response at each time point corresponds to a pair of retinal and eye velocities (e.g. *t*_1_ and *t*_2_ in near and far depths; pink and blue areas). **G**, Firing rates at each time bin are mapped to the retinal-eye velocity space according to the corresponding velocities. Responses in near-depth trials transform to lines with positive slopes (e.g. pink arrow and squares); response to far depths transform to lines with negative slopes (e.g. blue arrow and squares).

**Supplementary Figure 3.**
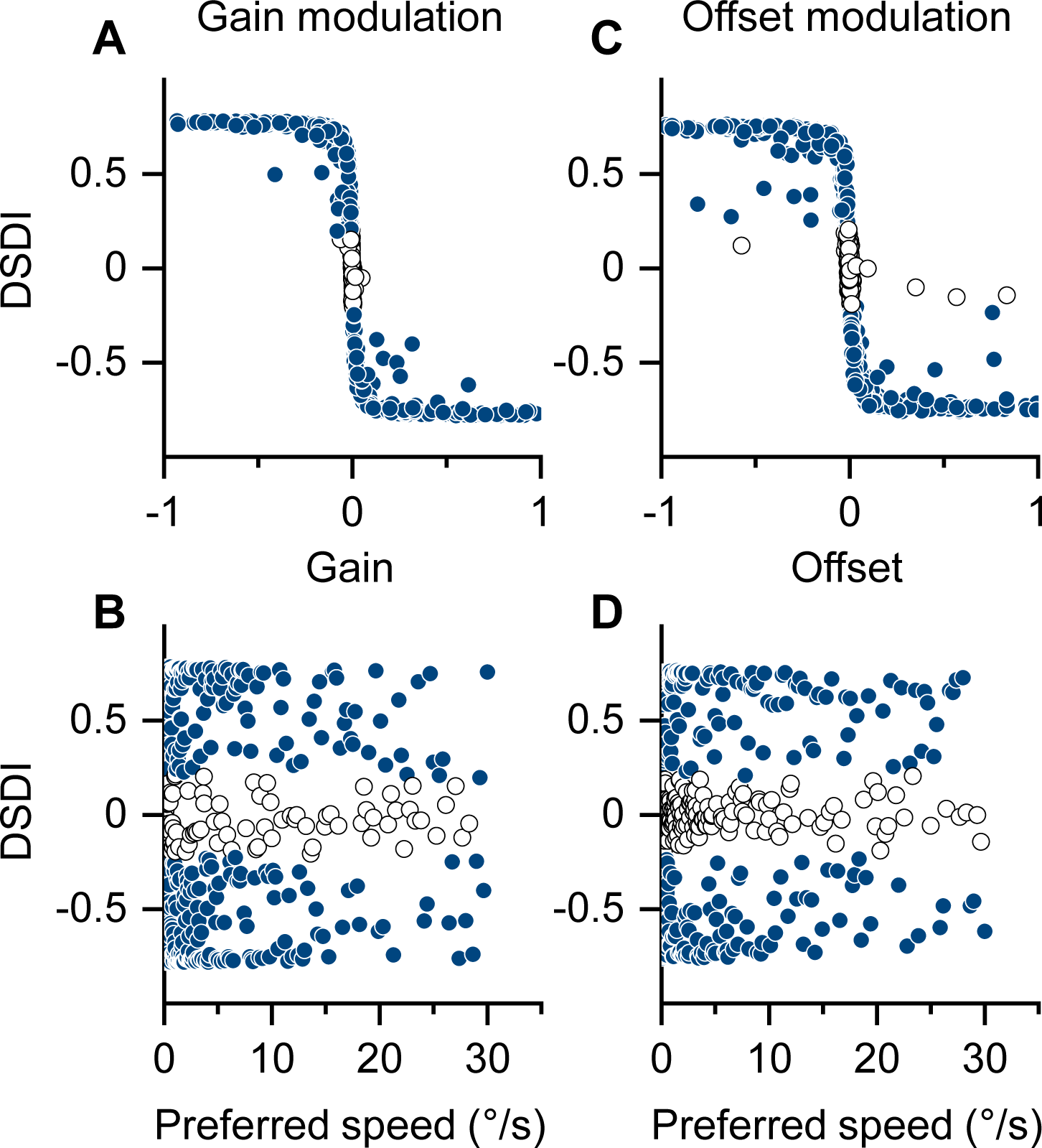
Simulating gain modulation and offset modulation. **A**, DSDI changes with the amount of multiplicative gain. **B**, Preferred speed of gain-modulated neurons does not predict the depth-sign selectivity. **C**, DSDI as a function of offset modulation. **D**, Preferred speed of offset-modulated neurons does not predict DSDI.

**Supplementary Figure 4.**
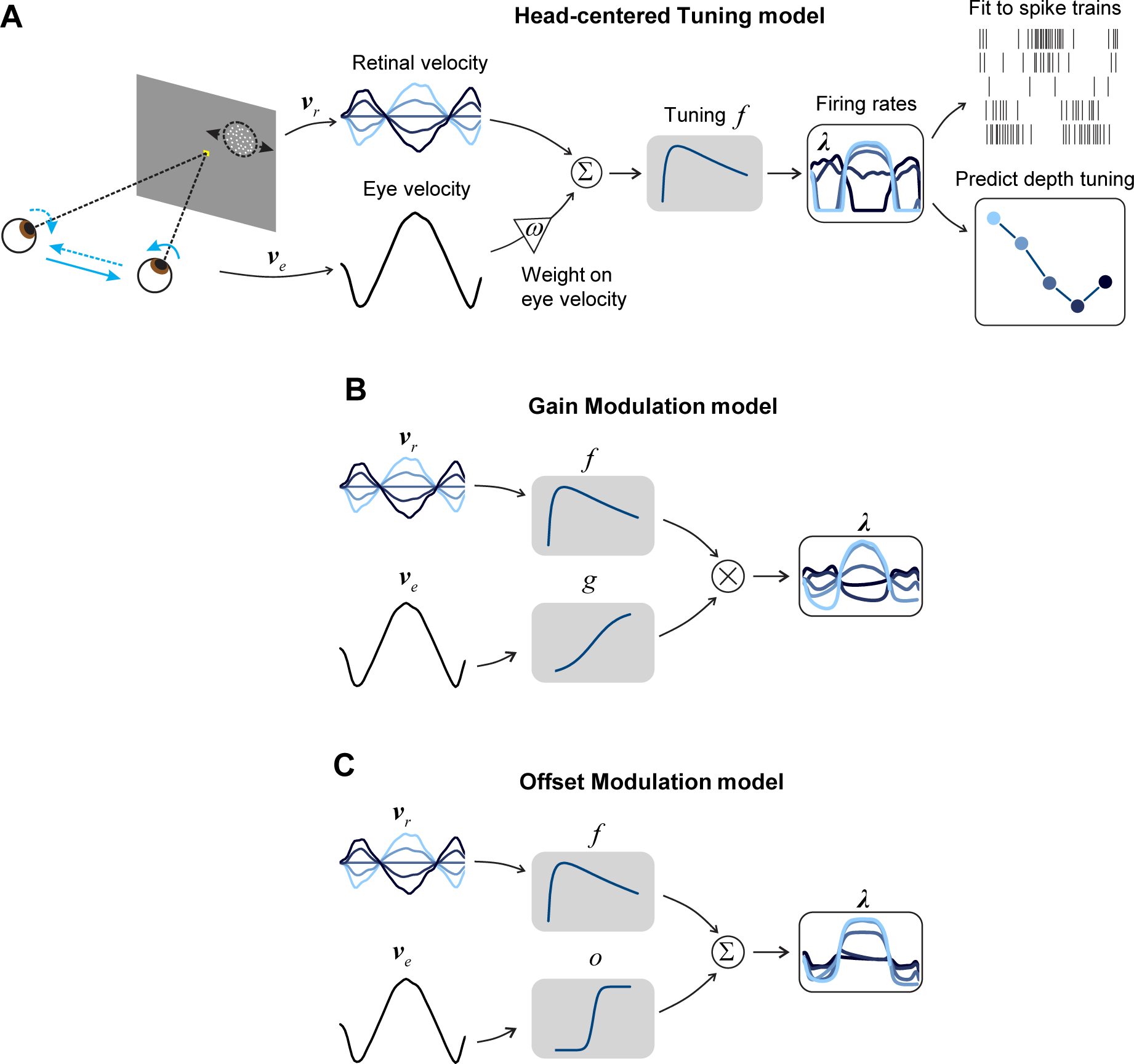
Illustrations of computational models. **A**, Scheme of model fitting on the head-center tuning (HT) model. Retinal and eye velocities, ***v****_r_* and ***v****_e_*, are linearly combined with a weight *ω* on eye velocity, and fed into a tuning function, *f* . The output firing rates, ***λ***, are then fit to spike trains to estimate model parameters. After fitting, the modeled response is used to predict depth tuning curve. **B**, Gain modulation model. Only retinal velocity is fed to the tuning function, while eye velocity multiplicatively modulates the tuning response by a gain function, *g*. **C**, Offset modulation model is similar with the GM model, except that eye velocity modulates the response in an additive fashion by an offset function, *o*. All models are fit to spike trains and are used to predict depth tuning curves.

**Supplementary Figure 5.**
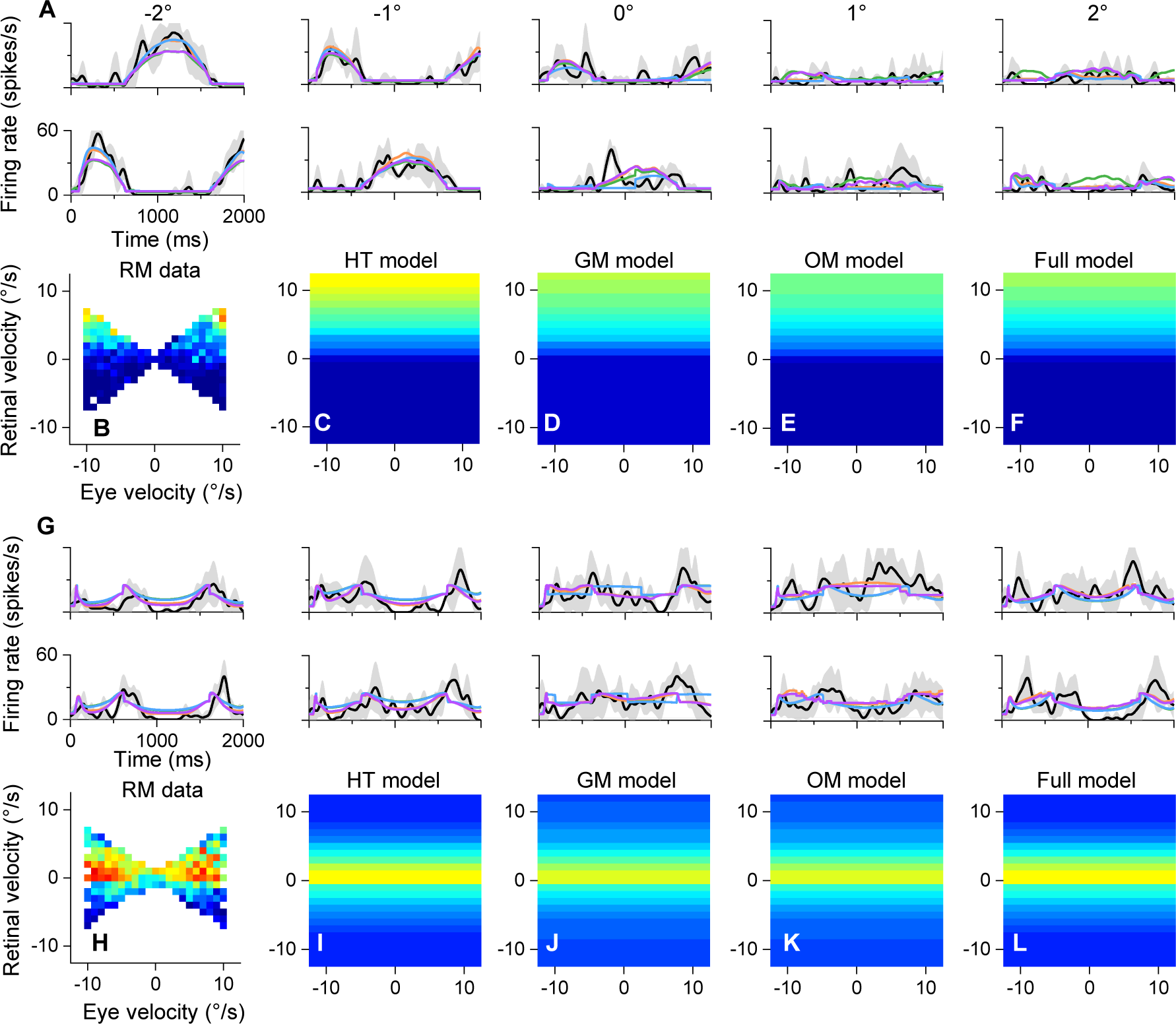
Model fitting results for PSTHs in the MP condition and joint tuning in the RM condition. **A-F**, Results for the same neuron shown in Fig. 3A-C and Fig. 6A-J. **A**, PSTHs and model fits in each depth and movement phases in the MP condition. First row shows the results for trials at 0° movement phase and the second row shows results for trials at 180° movement phase. Black curves show the PSTHs averaged across trials, gray shading indicates 1 SD around the mean. Colored curves show the model fits. **B**, joint tuning in the RM condition. **C-F**, Model predictions of the joint tuning in RM condition. **G-L**, Results for another example neuron shown in Fig. 3D-F and Fig. 6K-T.

**Supplementary Figure 6.**
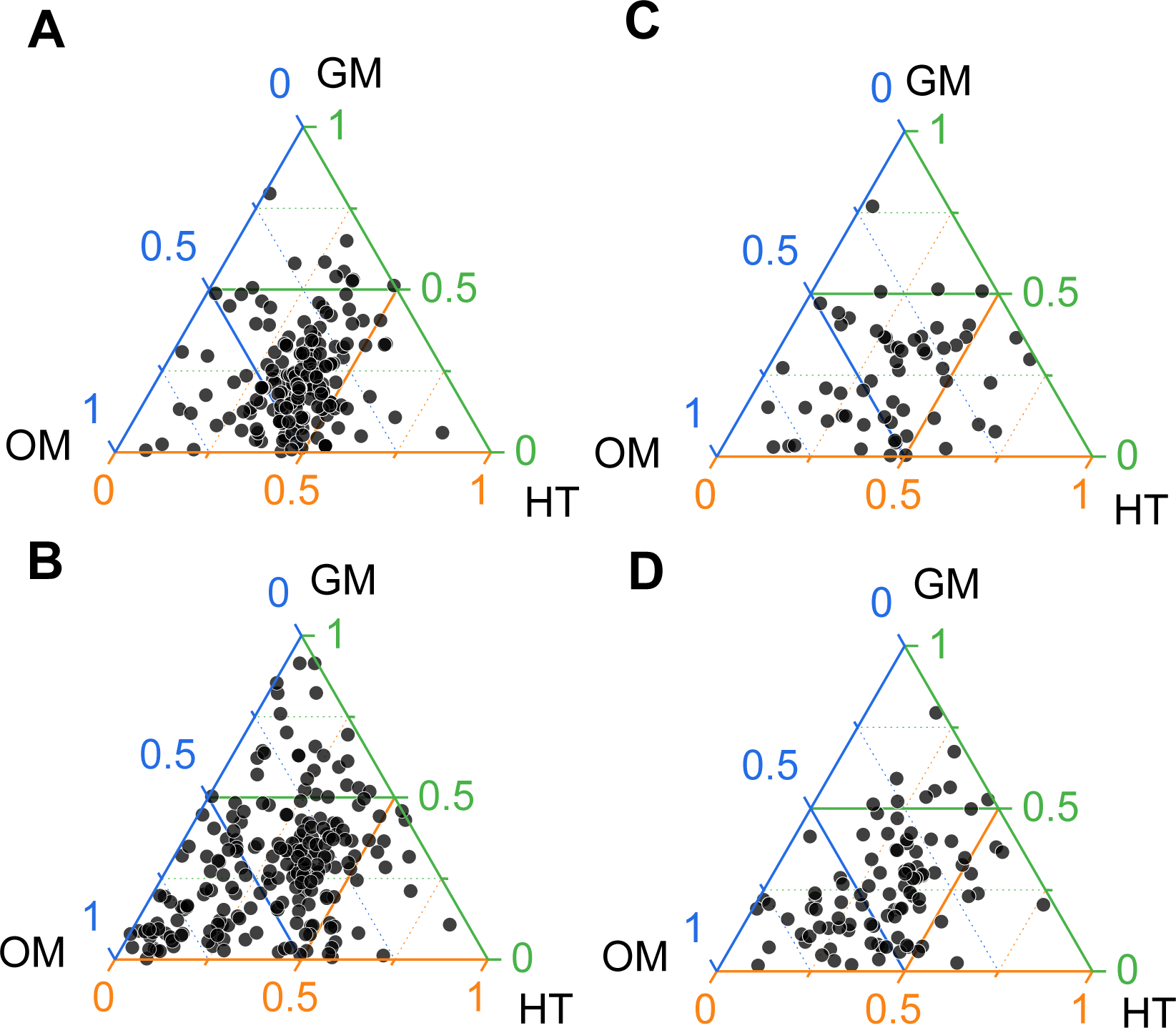
Relative contributions on BIC and DSDIs by each model component. **A**, Proportion of BIC change by adding one of the three model components to the control model in the MP condition. Dots closer to one of the vertices show a greater contribution of that model component. **B**, Proportion of |ΔDSDI| in three model components in the MP condition. Dots away from one of the vertices indicate that model predicted DSDI more accurately. **C**, Proportion of BIC change for each model in the DP condition. **D**, Proportion of |ΔDSDI| for each model in the DP condition.

**Supplementary Figure 7.**
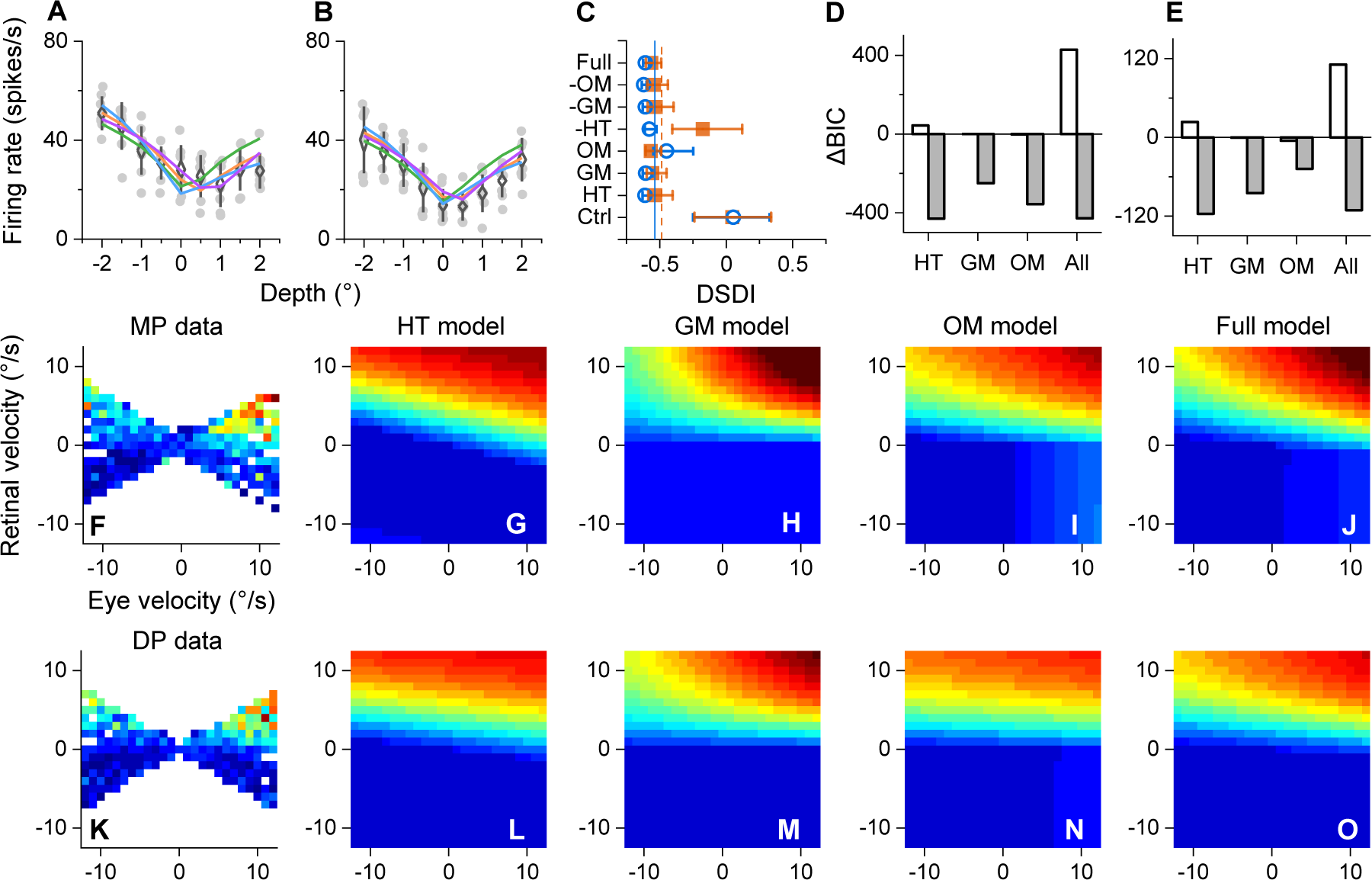
Additional example of model fitting in the DP condition. **A** and **B**, depth tuning curves in the MP and DP conditions, respectively. **C**, Predictions of DSDIs. **D** and **E**, Change of BIC by addition (gray bars) and removal (white bars) for each component. **F-J**, Joint tuning result (**F**) and model predictions (**G-J**) in the MP condition. **K-O**, results in the DP condition.

**Supplementary Figure 8.**
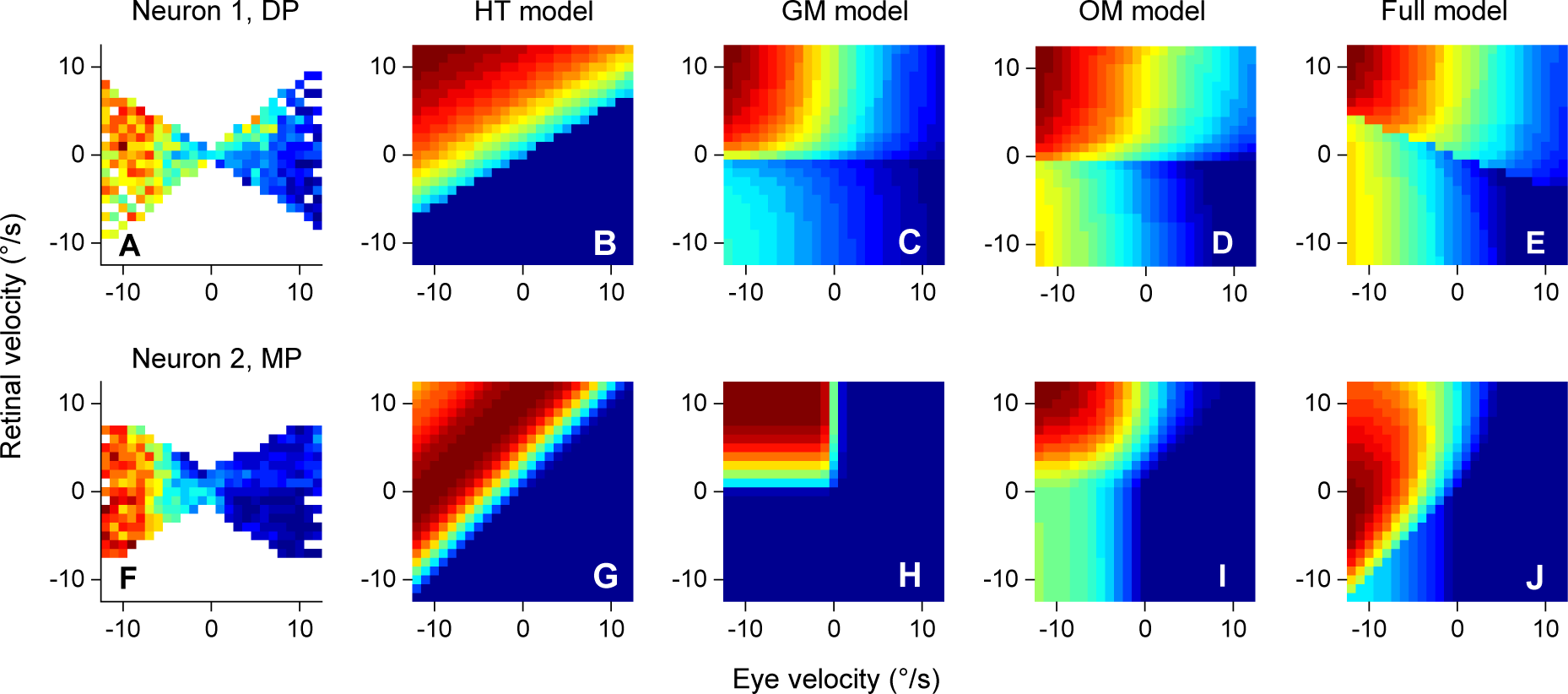
Joint tuning patterns dominated by eye velocity signal. **A**, Joint tuning profile of one example neuron that shows dominant modulation by eye velocity (x-axis) but not by retinal velocity (y-axis) in the DP condition. **B-E**, Model predictions of joint tuning pattern (**B**, HT model; **C**, GM model; **D**, OM model; **E**, full model). **F-J**, Another example neuron that shows dominant effect of eye velocity in the MP condition.

